# Taxonomic composition, community structure and molecular novelty of microeukaryotes in a temperate oligomesotrophic lake as revealed by metabarcoding

**DOI:** 10.1101/2022.06.30.498108

**Authors:** Konstantina Mitsi, Daniel J. Richter, Alicia S. Arroyo, David López-Escardó, Meritxell Antó, Antonio Guillén, Iñaki Ruiz-Trillo

**Affiliations:** Instituto de Biología Evolutiva (CSIC-Universitat Pompeu Fabra), Passeig Marítim de la Barceloneta, 37-49, 08003, Barcelona, Spain; Institut de Ciències del Mar (CSIC), Passeig Marítim de la Barceloneta, 37-49, 08003, Barcelona, Spain; I.E.S. “Escultor Daniel” C/ Gonzalo de Berceo, 49, 26005, Logroño, La Rioja, Spain; Departament de Genètica, Universitat de Barcelona, Avinguda Diagonal 643, 08028, Barcelona, Spain; Institució Catalana de recerca I Estudis Avançats (ICREA), Passeig Lluís Companys, 23, 08010, Barcelona, Spain; Departamento de Biodiversidad y Biología Evolutiva, Museo Nacional de Ciencias Naturales, Calle de José Gutiérrez Abascal, 2, 28006, Madrid, Spain

**Keywords:** microeukaryotes, 18S, V4, temperate, oligomesotrophic, protists, ASVs, novelty, phylogenetic placement, parasites

## Abstract

Microbial eukaryotes are diverse and ecologically important organisms, yet sampling constraints have hindered the understanding of their distribution and diversity in freshwater ecosystems. Metabarcoding has provided a powerful complement to traditional limnological studies, revealing an unprecedented diversity of protists in freshwater environments. Here, we aim to expand our knowledge of the ecology and diversity of protists in lacustrine ecosystems by targeting the V4 hypervariable region of the 18S rRNA gene in water column, sediment and biofilm samples collected from Sanabria Lake (Spain) and surrounding freshwater ecosystems. Sanabria is a temperate lake, which are relatively understudied by metabarcoding in comparison to alpine and polar lakes. The phylogenetic diversity of microbial eukaryotes detected in Sanabria span all currently recognized eukaryotic supergroups, with Stramenopiles being the most abundant and diverse supergroup in all sampling sites. Parasitic microeukaryotes account for 21% of the total protist ASVs identified in our study and were dominated by Chytridiomycota, both in terms of richness and abundance in all sampling sites. Sediments, biofilms and water column samples harbour distinct microbial communities. Phylogenetic placement of poorly assigned and abundant ASVs indicates molecular novelty inside Rhodophyta, Bigyra, early-branching Nucletmycea and Apusomonadida. In addition, we report the first freshwater incidence of the previously exclusively marine genera *Abeoforma* and *Sphaeroforma*. Our results contribute to a deeper understanding of microeukaryotic communities in freshwater ecosystems, and provide the first molecular reference for future biomonitoring surveys in Sanabria Lake.

## Introduction

The tree of eukaryotes is an ideal playground for biodiversity explorers. Although land plants, animals and fungi initially attracted most of researchers’ attention, the advent of molecular techniques in biodiversity assessment has revealed an enormous diversity of microbial eukaryotes outside these three groups (Pawlowski et al. 2012; del Campo et al. 2014). The paraphyletic assemblage of microbial eukaryotes is collectively referred to as protists (Archibald, Simpson, and Slamovits 2017). Protists are valuable from an evolutionary perspective because by studying their life traits we gain insights into the evolutionary processes that shaped the extant eukaryotic tree of life (Lang et al. 2002; del Campo et al. 2016; Grau-Bové et al. 2017; Gawryluk et al. 2019; Gabr, Grossman, and Bhattacharya 2020). In addition, protists are abundant, diverse and widespread organisms with key roles in important ecosystemic functions (Gao et al. 2019; Caron 2009; Gooday et al. 2020). However, despite their importance in different ecosystems as producers (Stoecker et al. 2009), grazers (Strom et al. 1997; W. D. Orsi et al. 2018), predators (Corno and Jürgens 2006) and parasites (Mahé et al. 2017), they have attracted less attention in comparison to their prokaryotic counterparts in environmental surveys (Caron et al. 2009).

Sampling based on morphological identification combined with environmental DNA (eDNA) analyses (Ruppert, Kline, and Rahman 2019) have shown that protists are everywhere (Epstein and López-García 2008). However, they are not everywhere equally studied. Microbial eukaryotes have been widely explored in marine ecosystems (P. López-García et al. 2001; Lovejoy, Massana, and Pedrós-Alió 2006; Worden, Cuvelier, and Bartlett 2006; Countway et al. 2007; Massana and Pedrós-Alió 2008; Alexander et al. 2009; Stoeck et al. 2010; Logares et al. 2014; Vargas et al. 2015), whereas there are fewer studies available regarding their distribution and diversity in soils (Fell et al. 2006; Shen et al. 2014; Moon-van der Staay et al. 2006; Mahé et al. 2017) and in freshwater systems (Šlapeta, Moreira, and López-García 2005; Debroas et al. 2017). Freshwater ecosystems are more fragmented and isolated (Dodson 1992; Reche et al. 2005) in comparison to the open ocean where microbial communities are transported by currents on a global scale (Villarino et al. 2018; Richter et al. 2020). This intrinsic lower connectivity of freshwater ecosystems hinders the dispersal of freshwater organisms and increases the genetic diversity (Dias et al. 2013).

Among freshwater habitats, lakes are undoubtedly the environments with the greatest number of molecular studies available (Charvet et al. 2012; Lepère et al. 2016; Boenigk et al. 2018). High mountain lakes (Kammerlander et al. 2015; Filker et al. 2016; Boenigk et al. 2018; Stoeck et al. 2014) and polar lakes (Daniel et al. 2016; Stoof-Leichsenring et al. 2020) have been extensively studied due to their extreme conditions of temperature, nutrient availability and UV radiation. These systems have been shown to harbour a high proportion of unclassified sequences within numerous eukaryotic lineages. Fewer molecular biodiversity surveys, however, have been conducted in lakes of temperate areas (Lefranc et al. 2005; Lepère et al. 2013).

In this study, we explore the diversity of microbial eukaryotes in Sanabria Lake (Spain), a temperate lake at an altitude of 1004.1 m above sea level. Sanabria is an oligotrophic to oligomesotrophic, warm, monomictic lake with winter circulation and summer stratification. In comparison to lakes of other trophic states, oligomesotrophic lakes harbour the richest and most diverse lentic organismal communities (Lefranc et al. 2005). Sanabria Lake is the biggest natural glacial lake in the Iberian Peninsula (Vega et al. 2005) and has already been the subject of many traditional limnological studies (Taboada 1913; Margalef 1955; Aldasoro et al., 1992; Vega et al., 1992; De Hoyos, 1996; De Hoyos et al., 1998; De Hoyos and Comín, 1999; Negro, De Hoyos, and Aldasoro 2003; Luque 2002; Jambrina-Enríquez et al. 2014; Pahissa, Fernández-Enríquez, and De Hoyos 2015; Llorente and Seoane 2020) Negro et al., 2000; Luque & Julià, 2002; Rico et al., 2007; Jambrina-Enríquez et al., 2014; Pahissa et al., 2015; Llorente and Seoane 2020). However, no molecular data are currently available for this freshwater system.

The aim of this study is to explore the eukaryotic microbial community of Sanabria Lake using a massively parallel sequencing approach. To do so, we targeted the V4 hypervariable region of the 18S rDNA gene. We explored the taxonomic composition of the microbial eukaryotes present in the lake and the surrounding freshwater systems, including an exploration of the protist parasite diversity. We also assessed the community structure and the compositional heterogeneity across different sampling sites, habitats and filter size fractions. Finally, we analysed our samples using a phylogenetic placement approach to quantify molecular novelty and we identified the branches of the eukaryotic tree that harbour potentially novel abundant taxa. Sanabria Lake is a protected biotope and it is under continuous monitoring. This is the first biodiversity study of Sanabria Lake that is based on molecular data, which will constitute a reference for future monitoring efforts.

## Materials and Methods

### Study area

Sanabria Lake is situated in the NW of Spain (42 7’ 30’’ N, 06 3’00”W) between the provinces of León and Zamora at an altitude of 1004.1 m above sea level. It was formed by glacial erosion after the Würm glaciation in the Pleistocene, and it is the only lake formed by a terminal moraine in the Iberian Peninsula (Margalef 1983). Sanabria Lake belongs to the Duero River Basin that has a total drainage surface of 127.3 km^2^ (De Hoyos, 1996) and its main tributary is the Tera River. The surface of the lake is 3.46 km^2^ (De Hoyos, 1996). It is divided longitudinally into two basins, one in the west with maximum depth of 46 m and another in the east with maximum depth of 51 m (Vega et al. 2005). The shoreline length is 9518 m and the maximum width is observed in the eastern basin (1530 m) (Vega et al. 2005). Regarding its mixing characteristics, Sanabria Lake is a warm, monomictic, holomictic lake (Vega et al 1992). The mixing period extends from late November to early March, when a thermocline normally appears (Vega et al 1992). No anoxic conditions have been observed in any layer of the water column during the thermal stratification (Vega et al 1992; de Hoyos 1996; Pahissa, Fernández-Enríquez, and De Hoyos 2015). Sanabria Lake is considered as oligotrophic to oligomesotrophic in view of its low levels of chlorophyll *a*, nutrient concentration, phytoplanktonic biovolume values and production rates (Planas 1991; Vega et al 1992; de Hoyos 1996; Pahissa, Fernández-Enríquez, and De Hoyos 2015; Llorente and Seoane 2020; De Hoyos and Comín 1999). The oligotrophic state of the lake is a result of its geology. Its drainage basin runs over an acid rock substrate (gneiss and granodiorites) of low solubility, making the water very poor in salts (Rico et al 2007). The lake is part of the Sanabria Lake Natural Park (BOE 1978), a protected area that supports a population of 2 small villages (∼ 200 residents), one in the north and the other in the west side of the lake. During the summer, the National Park receives a high influx of tourists and there are three camping sites, all located on the east side of the lake. Since 2012, the Duero International Biological station has raised concerns that Sanabria Lake is undergoing a eutrophication process due to contamination from a deficient sewage depuration system (Guillen, 2015). However, studies based on pigment measurements and microscopy observation of the phytoplankton community do not support the eutrophication scenario and confirm the current oligotrophic state of the lake (Pahissa, Fernández-Enríquez, and De Hoyos 2015; Llorente and Seoane 2020).

### Sampling

We sampled in March 2016 at the beginning of the thermal stratification. We chose this time point because the physicochemical conditions of the lake are homogeneous after the winter mixing period. We expected that microbial eukaryotes would be homogeneously distributed across the lake and that this even distribution would increase sampling efficiency. In addition, this is the time of the year with the least anthropogenic impact. Thus, any disturbance detected would indicate a permanent change rather than a temporal variation due to the presence of a stressor.

To explore the diversity and the relative abundance of microbial eukaryotes in Sanabria Lake, we collected 82 samples of water, sediment and biofilms from ten different locations. Water samples were collected from five sampling sites in the lake basin (S1-S5), a tributary stream (S6-S7) and a nearby pond (S8-S10) (Figure 1, Supplementary Table 1). The sampling was designed to include as many different habitats as possible: i) two pelagic sampling sites in the lake, one in the deepest point of the west basin (S1) and another in the deepest point of the east basin (S4), ii) the mouth of River Tera (S2), a sampling site experiencing the greatest anthropogenic impact in the studied ecosystem, iii) a coastal area in the lake near two islets (Islas Moras) (S3), iv) sulphurous waters in the east basin (S5), v) water from a tributary stream (S6) and vi) water from three nearshore sites in a nearby pond (S8-S10). For each of the sampling sites S1-S5, we collected between 0.5 and 1 litres of water from the surface, the Deep Chlorophyll Maximum (DCM) and the deepest point above the sediment. We measured standard variables (turbidity, temperature, fluorescence) in the sampling sites of the lake’s main water body (S1-S5) using a CTD SD204 (SAIV A/S) device. We prefiltered the water samples using filters of 2000 μm and 200 μm to remove debris and large multicellular organisms. Size fractions above 200 μm were discarded and not included in the study. Subsequently, we filtered sequentially using filters of 20 μm, 5 μm and 0.8 μm targeting the microplankton (20-200 μm), the nanoplankton (5-20 μm) and the pico/nanoplankton (0.8-5 μm) communities respectively. We additionally collected sediments from three sampling sites (S2, S3, S7) and 12 epilithic biofilm samples from one sampling site (S3). All samples were placed in 2ml cryovials and were stored at -80 °C prior to DNA extraction.

**Figure 1.**
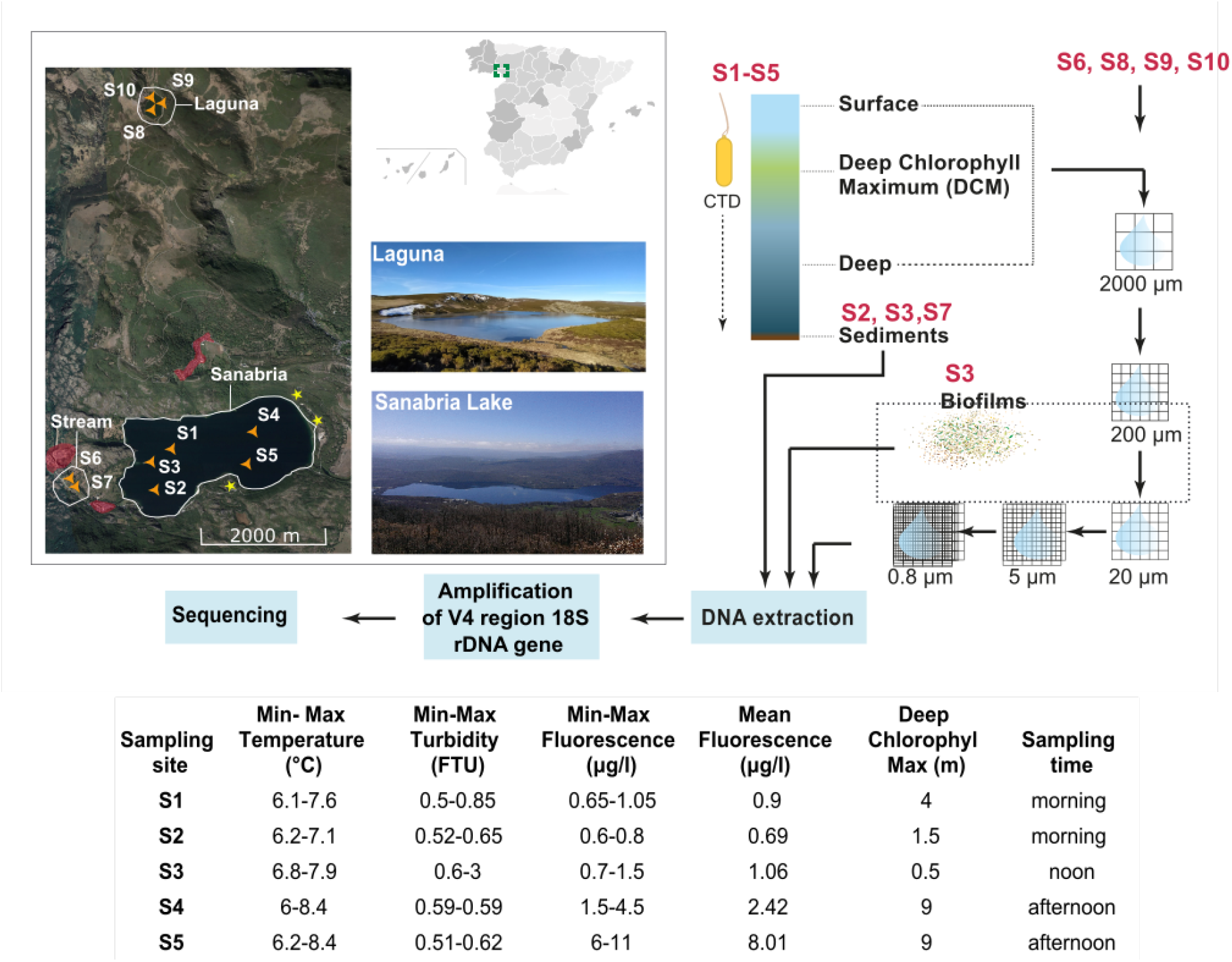
Sampling information. The grey map on the top right of the box shows the position of Sanabria Lake in the Iberian Peninsula. The map on the left shows the sampling sites pointed by orange triangles. Nearby villages are delimited by red coloured areas. Camping sites are pointed by yellow stars. The sampling protocol is detailed in the upper right part of the figure. Filters of 2000μm and 200μm contained mainly multicellular organisms and they were not sequenced. Sample S6 is water and S7 is sediment from an upstream tributary stream. Samples S8-S10 are water samples from a nearby small pond (Laguna de los Peces) that is not connected to the main water body. In the table, we present CTD data collected in Sanabria Lake (sites S1-S5).

### DNA extraction, PCR amplification and sequencing

We chopped the filters using sterile scissors and we homogenised the sediments and the biofilms before further processing. We extracted whole genomic DNA using the standard protocol of the PowerSoil DNA Isolation Kit (MO BIO). We amplified the V4 hypervariable region of the 18S rRNA gene using the universal eukaryotic V4 primers TAReuk454FWD1 (5’-CCAGCA(G⁄C)C(C⁄T)GCGGTAATTCC-3’) and TAReukREV3 (5’-ACTTTCGTTCTTGAT(C⁄T)(A⁄G)A-3’) (Stoeck et al. 2010). The amplicons were sequenced using the Illumina MiSeq platform at Centro Nacional de Análisis Genómico (CNAG, Barcelona, Spain). The sequences were demultiplexed and the barcodes were trimmed by the sequencing facility. The raw sequencing data were submitted to the European Nucleotide Archive (ENA) under the accession number PRJEB23911 (ERP105690).

### Amplicon Sequence Variants (ASVs) analysis

We analysed the raw reads following a clustering-free approach. We processed the Illumina demultiplexed paired-end sequenced dataset using the R package dada2 (Callahan et al. 2016). We visualised the read quality profiles using the function plotQualityProfile. The quality of the forward and reverse reads started decreasing after the position 250 and 240, respectively. We used the function filterAndTrim to discard low quality sequences using standard filtering parameters (maxN = 0, maxEE = c(2,2), truncQ = 2, rm.phix = TRUE, compress = FALSE, multithread = TRUE) and to trim the forward reads in the position 250 and the reverse reads in the position 240. We calculated the error model from our data with the function learnErrors and we visualised the estimated error rates with the function plotErrors. We then dereplicated the reads using the function derepFastq and we inferred sequence-variants from our dereplicated data using the function dada. With the function mergePairs we merged the forward and the reverse reads to obtain the full denoised sequences using the default 12 bases overlap region and we removed the paired reads that did not exactly overlap. We constructed our amplicon sequence variant table (ASV) table using makeSequenceTable, then removed chimeras with removeBimeraDenovo and finally inspected the number of reads that made it through each step of our analysis (Supplementary Table 2). We assigned taxonomy with the function assignTaxonomy that uses the Ribosomal Database Project (RDP) classifier together with the Protist Ribosomal Reference database (PR2) (v. 4.12.0) (Guillou et al. 2013) formatted for DADA2 (https://github.com/pr2database/pr2database/releases). We generated an ASVs table that contains a total of 31225 ASVs, the ASV counts per sample and their taxonomy using Biostrings::DNAStringSet and Biostrings::writeXStringSet functions.

### Statistical analyses

We combined the taxonomy, the abundance and the metadata to generate a phyloseq object (McMurdie and Holmes 2013) (Supplementary Table 3). We generated the different phyloseq datasets used in this study by subsetting (phyloseq::subset_samples, phyloseq::subset_taxa) the initial phyloseq object (Supplementary Table 3). We plotted rarefaction curves using the function phyloseq::rarecurve to explore whether all included samples have reached saturation. Samples that did not reach saturation were removed. We used the phyloseq (McMurdie and Holmes 2013), psadd (https://rdrr.io/github/cpauvert/psadd/) and Fantaxtic (https://rdrr.io/github/gmteunisse/Fantaxtic/) R packages to plot the taxonomic composition of the datasets.

We rarefied (phyloseq::rarefy_even_depth) each dataset at the minimum sample depth (Supplementary Table 3) in order to simulate even numbers of reads per sample. Rarefaction enables clustering samples according to their biological origin and permits fair comparison of diversity metrics among the samples (Weiss et al. 2017). We calculated alpha (Supplementary Table 4) and beta diversity in the subsampled datasets. Alpha diversity metrics (Observed species, Chao1, se.chao1, ACE, se.ACE, Shannon, Simpson, InvSimpson, Fisher) were calculated using the function phyloseq::estimate_richness. The significance of the difference in species richness was tested with pairwise comparisons using the Wilcoxon rank sum test, controlling for family wise error rate with the Holm–Bonferroni method (Xie et al. 2017). Evenness was calculated according to Pielou (Pielou 1966) and plotted as violin plots in the ggplot2 R package (Wickham 2009). Beta diversity was measured using the Bray–Curtis dissimilarity statistic after converting ASVs abundances to frequencies within samples. To test the effects of habitat, sampling depth, sampling site, chlorophyll maximum and thermocline across samples, we conducted permutational multivariate analyses of variance (PERMANOVA) based on Wisconsin-standardized Bray-Curtis dissimilarities (Supplementary Table 6), using the adonis function of the vegan package. Patterns of beta diversity were assessed via non-metric multidimensional scaling ordination (NMDS) also on Bray-Curtis dissimilarities using the function phyloseq::ordinate and were plotted using the function phyloseq::plot_ordination. To test the significance of groups revealed by NMDS, we applied analysis of similarity (ANOSIM) tests with 999 permutations (Supplementary Table 7).

### Phylogenetic analysis

Reference trees were built in RAxML (Stamatakis 2014) under the GTRGAMMA model. Node support was estimated by 100 rapid bootstrap replicates. Phylogenetic placements were produced using RAxML-EPA (Berger and Stamatakis 2011) and visualised with iTOL (Letunic and Bork 2011). Each query was placed in multiple branches until the total like_weight_ratio summed to 1. All alignments were built with MAFFT v7.309 (Katoh and Standley 2013) with automatically selected strategy according to data size and were trimmed with trimal v1.4.rev15 (Capella-Gutiérrez, Silla-Martínez, and Gabaldón 2009) using the -automated1 algorithm.

## Results and Discussion

### Abiotic parameters indicate ecological disturbance in the east basin of Sanabria Lake

Sanabria Lake is a warm monomictic lake with water circulation during winter and thermal stratification that begins in the spring and continues until the end of the summer (Vega et al 1992). Samples were collected at the beginning of the thermal stratification period when the eukaryotic community is expected to be homogeneously distributed in the water body following the winter mixing. This period also coincides with the time of the year with the lowest direct anthropogenic impact. Water temperature at the surface ranged from 7.1 °C to 8.4 °C and in the deepest sampling points ranged from 6 °C to 6.85 °C, with a mean range of 1.62 °C (Figure 1, Supplementary Table 1). These temperature measurements agree with the previously recorded temperatures during the homeothermic state of the lake that range between 4 to 7 °C (Vega et al 1992) and confirm the mixing state of the lake.

We assessed the trophic state of Sanabria Lake based on water turbidity and chlorophyll *a* values. Water turbidity is measured in FTU (Formazin Turbidity Units) and is an indicator of the trophic state of a lake as it is related to the concentration, type and size of the suspended particles in the water (Çako, Baci, and Shena 2013). During our sampling, turbidity values in Sanabria Lake were extremely low in all the sampling sites and ranged from 0.5 to 0.85 FTU (Figure 1). These values are comparable to those in ultra-oligotrophic alpine lakes (Chanudet and Filella 2007). Chlorophyll *a* is a reliable indicator to assess the trophic state of a lake with high values to correspond to more eutrophic ecosystems (Poikane et al. 2014). Chlorophyll *a* mean values in Sanabria Lake have increased in the last fifty years (Supplementary Table 5) but they have not exceeded the levels that characterise oligotrophic lacustrine ecosystems. Together, these measurements confirm the overall oligotrophic status of the lake at the time of sampling.

We observed that chlorophyll *a* values differed between east and west basin during our sampling (Figure 1). In Sanabriás west basin (samples S1-S3), the mean value of chlorophyll *a* was below the reference value (1.5 μg/L). The reference value defines the equilibrium ecological state of the lake and confirms the absence of ecological disturbances. However, the mean values of chlorophyll *a* in Sanabriás east basin (samples S4-S5) exceeded the reference values indicating the presence of ecological disturbance (Figure 1). Values of chlorophyll *a* above 4.2 μg/L are linked to a Good-Moderate ecological state and values above 7.1 μg/L are linked to a Moderate-Poor ecological state (BOE, 2015). Our results showed that there was some ecological disturbance that altered the values of chlorophyll *a* in the east basin of Sanabria Lake at the time of sampling. The altered values of chlorophyll *a* in the east basin may be related to higher anthropogenic impact due to the presence of three camping sites on this side of the lake. Chlorophyll *a* values measured in Sanabriás east basin in March 2017 (Llorente and Seoane 2020) are lower than the ones presented in our study, implying that the stressor was temporal and that water quality has been restored.

### The V4 hypervariable region captures the microeukaryotic diversity of Sanabria Lake

To characterise the diversity of the eukaryotic community in Sanabria Lake, we sequenced the V4 hypervariable region of the 18S small subunit (SSU) rRNA gene. We chose to sequence the V4 over other hypervariable regions of the 18S rRNA gene because it provides taxonomic resolution close to that of the full-length gene (Dunthorn et al. 2012; Hu et al. 2015) and it is the most suitable hypervariable region to use for phylogenetic placement (Mahé et al. 2017). A total of 15,947,744 reads from 82 samples were filtered, dereplicated and merged resulting in 31,225 Amplicon Sequencing Variants (ASVs). The study of multicellular organisms was out of the scope of the present work and thus most multicellular organisms were discarded by using physical filters of 2000 μm and 200 μm. However, some environmental DNA (eDNA) that originates from cellular material shed by multicellular organisms into the lake was sequenced together with the community DNA of unicellular eukaryotes. For our subsequent analyses, we bioinformatically filtered out all ASVs that were assigned to animals (Division/Class = Metazoa), land plants (Division = Streptophyta) and typical terrestrial fungi (Class/Order = Ascomycota, Class/Order = Basidiomycota) (Supplementary Table 3, dataset D3). After the removal of multicellular taxa, 27,790 microeukaryotic (protist) ASVs remained. We evaluated the sampling depth and the representation of microbial eukaryotes in our samples using rarefaction curves (Supplementary Figure 1). The curves reached a plateau for all samples, indicating that most of the microbial richness present in Sanabria Lake and the surrounding freshwater systems was successfully captured by our study.

### Spatial biodiversity patterns

To evaluate the intra-sample diversity of Sanabria Lake and the surrounding water bodies, we calculated nine different alpha-diversity indices (Supplementary Table 4). To avoid potential biases in diversity estimates due to differences in the total number of reads, we randomly subsampled the ASVs to the minimum depth of our dataset (Supplementary Table 3, dataset D3, min sample depth = 31361 reads) before calculating the alpha-diversity indices. The number of total taxa reported was not affected by subsampling. We compared the diversity of the different water body types and we found that samples collected in the tributary stream showed significantly higher intra-sample diversity (Figure 2) and greater evenness (Supplementary Figure 10) compared to samples from Laguna (pond) and Sanabria (lake) (Wilcoxon rank sum test P value<0.01). Previous studies have shown that small water bodies like ponds and streams can contribute significantly to regional biodiversity of macrophytes and macroinvertebrates (Williams et al. 2004). Our data support the hypothesis that the same is true for microeukaryotes. This result pinpoints the importance of small water bodies as biodiversity reservoirs and contrasts with their relative status in national monitoring and protection strategies, where they are frequently ignored. Regarding the different habitats, sediments harbour the richest microeukaryotic communities (Figure 2). Sediments have been shown to harbour richer communities than the water column for other groups of organisms like bacteria (Eckert et al. 2020) and marine diatoms (Piredda et al. 2018). However, we cannot exclude that part of the diversity recorded in the sediments can be attributed to either dormant stages of planktonic microeukaryotes or dead cells that were recently settled from the water column.

**Figure 2.**
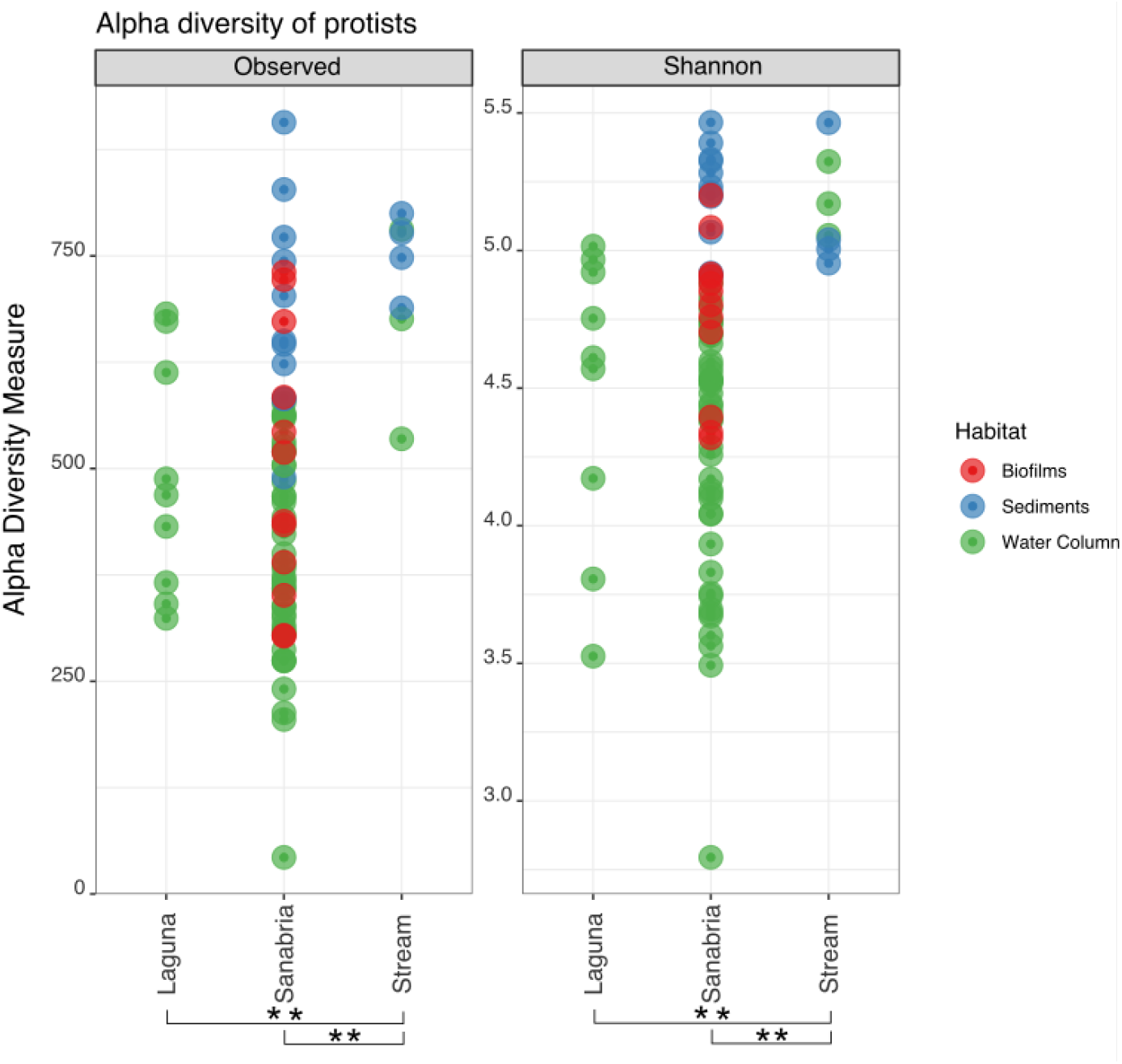
Alpha diversity of protists across the sampling sites. Each dot represents a sample and the colour code indicates the habitat of origin. Significant differences between pairs are indicated by double asterisks (p-value ≤0.01 **).

To test the effect of abiotic factors in the protist community structure we observed, we carried out permutational multivariate analysis of variance (PERMANOVA) of Bray-Curtis dissimilarities of the ASVs between communities as a function of sample spatial origin (Supplementary Table 6). All factors tested by PERMANOVA tests revealed significant differences in protist communities as a function of site (Sanabria Lake, Laguna, Stream), sampling site (S1-S10), position regarding the chlorophyll maximum (on-off), position regarding the thermocline and habitat (water column, sediments, biofilms) (Supplementary Table 6).

To visualise the compositional differences in the community structure of protists we applied nonmetric multidimensional scaling (NMDS). The communities from Sanabria Lake, the tributary stream and the Laguna were clearly separated in an ordination based on sampling site (Figure 3A). The samples were also distributed along the first axis (NMDS1) as a function of the habitat, with water column samples occupying the first three quadrants, biofilm samples in the third and fourth quadrant and sediment samples occupying only the fourth quadrant (Figure 3A). The community of microbial eukaryotes in the water column of Sanabria Lake was clearly segregated as a function of the size fraction of the filter and not the sampling depth (Figure 3B). This is what we expected given that we sampled at the beginning of the thermal stratification after the winter mixing at the point of maximum homogeneity of the community. As we observed that chlorophyll *a* values differed between east (S1,S2, and S3 sampling sites) and west basin (S4 and S5 sampling sites) (Figure 1) we investigated using a NMDS plot whether the microeukaryotic communities of east and west basin are grouped together but we did not observe such grouping (Supplementary Figure 2)

**Figure 3.**
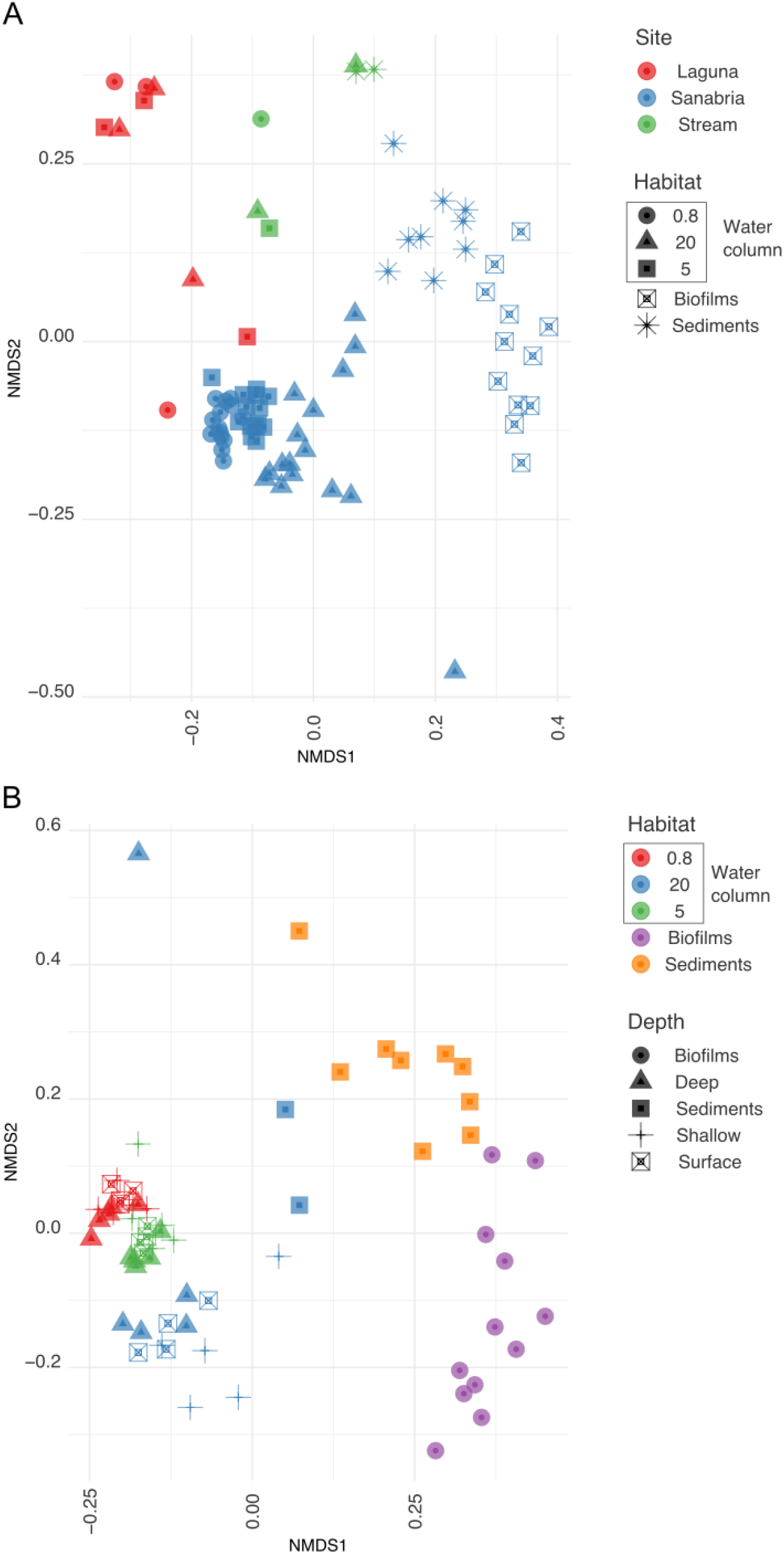
Reduced-space NMDS plot showing microbial eukaryotes community structure based on Bray–Curtis dissimilarity. A) Dissimilarity calculated from the rarefied at even depth (31361 reads) abundances of protist ASVs in all samples (dataset D3). The plot is divided in four quadrants horizontally and four quadrants vertically, delimited by grey lines. (Stress = 0.2083063, Procrustes: rmse 0.03612112 max resid 0.1863833), B) Dissimilarity calculated from the abundances of rarefied at even depth (31361 reads) ASVs present only in Sanabria samples (dataset D5). The plot is divided in three quadrants horizontally and five quadrants vertically, delimited by grey lines. (Stress = 0.1676406, Procrustes: rmse 2.784844e-06 max resid 1.93149e-05)

Our observations were statistically supported by ANOSIM tests (Supplementary Table 7), which showed significant and marked differences among communities according to habitat, sampling site and depth (Supplementary Table 1). Our results suggest that the community structure in Sanabria Lake and the surrounding freshwaters is characterised by spatial variation. The habitat is a major factor that shapes the community structure after the winter mixing period. Sediments, biofilms and water column harbour compositionally heterogeneous microbial communities that are driven by the unique environmental parameters that characterise them.

### Taxonomic composition of the protist community

To gain an overview of the microeukaryotic taxonomic composition in the Sanabria Lake and the surrounding freshwater systems, we plotted the relative abundance of ASVs at division level (based on the PR2 taxonomy) across sampling sites (Figure 4). The phylogenetic diversity of ASVs covered all currently recognized eukaryotic supergroups (Adl et al. 2019). The group of Stramenopiles was the most abundant supergroup in all sampling sites, accounting for the 33% of total reads in Sanabria Lake, 34% in the nearby pond (Laguna) and 40% in the tributary stream respectively (Supplementary Figures 3, 4, and 5). In addition to being abundant, Stramenopiles were diverse, encompassing 22% of total ASV richness (6,988 ASVs) (Supplementary Table 3). Among Stramenopiles, Ochrophyta was the most abundant group in all sampling sites (Supplementary Figure 6). Most Ochrophyta in the tributary stream (85%) and Laguna (81%) were affiliated with Chrysophyceae (Supplementary Figure 6), a group that is generally common in low-nutrient lakes (Nicholls and Wujek 2003). In Sanabria Lake, together with the Chrysophyceae (36%), we report a high relative abundance of Bacillariophyta (37%) and Synurophyceae (24%) within Ochrophyta, two phototrophic lineages that produce silica skeletons or scales (Supplementary Figure 6). Alveolata was the second most abundant and diverse supergroup, accounting for 26%-28% of the total eukaryotic reads in each site (Supplementary Figures 3, 4, and 5) and a total of 4,609 ASVs in the study (Supplementary Table 3).

**Figure 4.**
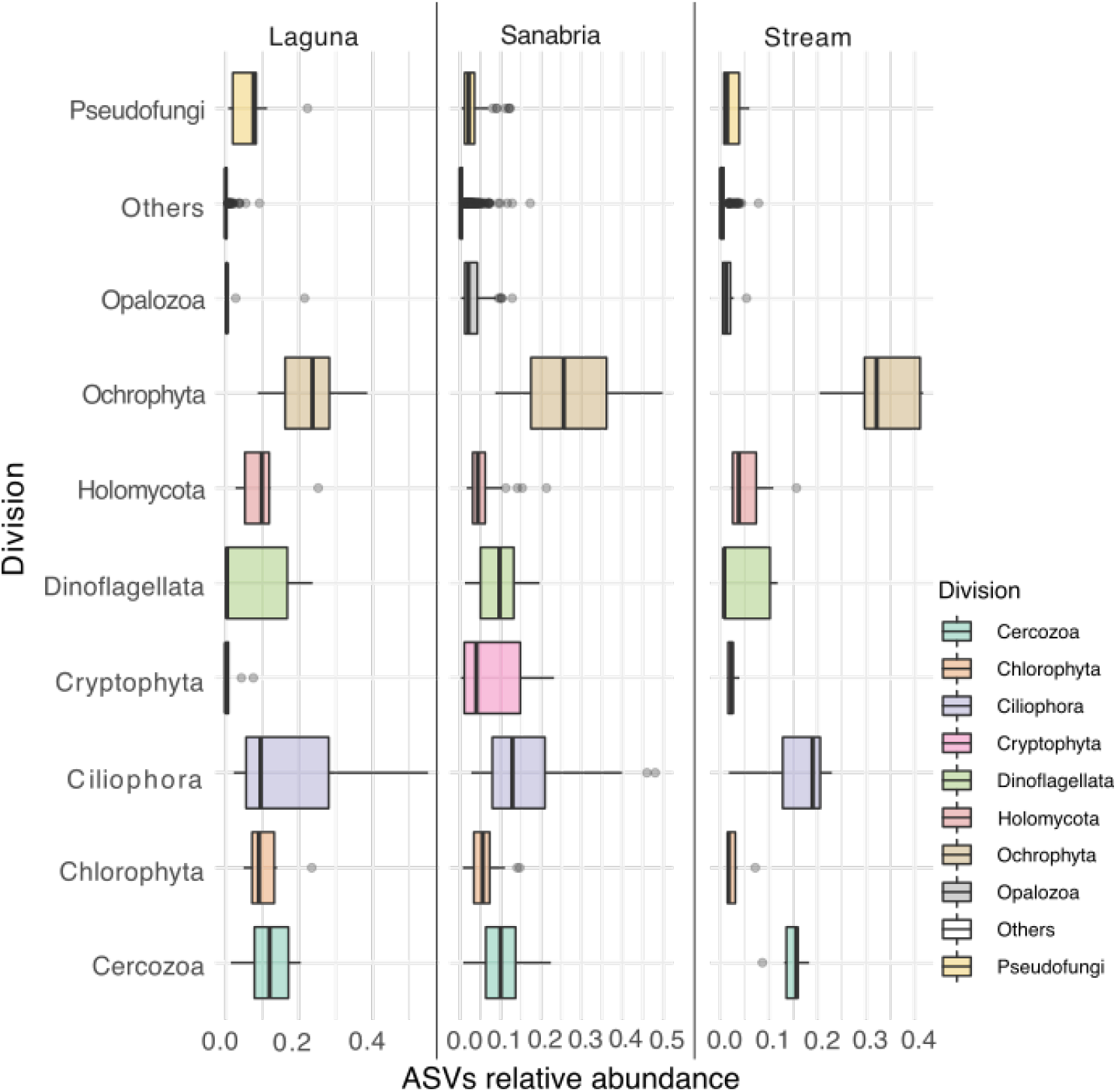
Distribution and relative abundance of the eukaryotic divisions across sampling sites as defined by ASVs. “Others” group together all taxa with relative abundance less than 1%. The boxes represent the interquartile range (IQR) between the first and third quartiles and the vertical line inside the box defines the median.

The plankton community of Sanabria Lake (excluding the surrounding freshwater systems) was dominated by Ochrophyta (in the Stramenopiles supergroup; 26%), Ciliophora (Alveolata; 14%), Dinoflagellata (Alveolata; 10%), Cercozoa (Rhizaria; 10%), Cryptophyta (10%) and unicellular Opisthokonta (7%) (Supplementary Figures 3). The presence of all these groups except for unicellular Opisthokonta was previously documented by light microscopy in Sanabria Lake (Vega et al. 1992). We further explored the taxonomic composition of Sanabria Lake by separately examining benthic and pelagic samples. The taxonomic composition of the benthic protist community, as represented by ASVs in the sediments, was dominated by Stramenopiles (36%), Alveolata (29%), Rhizaria (13%), Opisthokonta (7%), Amoebozoa (5%), Archaeplastida (3%), Hacrobia (3%) and Excavata (3%). In contrast, the planktonic microbial community was characterised by the prevalence of Hacrobia (16%) as the third most abundant eukaryotic supergroup after Stramenopiles (31%) and Alveolata (28%). The planktonic Hacrobia (Cavalier-Smith, Chao, and Lewis 2015) in Sanabria Lake included Cryptophyceae (84%), Katablepharidophyta (13%), Centroheliozoa (1.5%), Telonemia (1%) and Haptophyta (0.5%). Excluding Cryptophyceae, this is the first record for these taxonomic groups in Sanabria Lake. (Katablepharidophyta were previously classified inside Cryptophyceae until electron microscopy and molecular phylogenies provided evidence to consider them as a separate taxonomic group (Okamoto and Inouye 2005).)

### Protist parasites in a temperate oligomesotrophic lake

Here we provide the first description of the taxonomic composition of the unicellular eukaryotic parasites (Supplememtary Table 3, dataset D6) present in Sanabria Lake, the biggest natural lake in the Iberian Peninsula. Parasitic unicellular eukaryotes modulate large scale ecological processes by regulating the abundance and dynamics of their hosts population (Bråte et al. 2010). As their study by microscopy is tedious, little was known about their prevalence in freshwater systems until the advent of metabarcoding (Frenken et al. 2017).

The parasites accounted for 21.3% (5,925) of the total protist ASVs identified in our study. The parasitic community composition was dominated in all sampling sites by Chytridiomycota, whose relative abundance within parasitic taxa was 29% in the tributary stream, 32% in Sanabria Lake and 42% in Laguna. The prevalence of Chytridiomycota in the pelagic zone of lakes has been previously reported (Lefèvre et al. 2007; Sime-Ngando, Lefevre, and Gleason 2011). Chytridiomycota, which includes more than 1,000 described species (Powell 1993; Shearer et al. 2007), was also the most diverse group of parasites in our study, including more than 2,200 of the 5,925 total parasite ASVs, distributed among more than 50 genera (Supplementary Figure 7). Almost half of the chytrids in terms of abundance identified in our study belonged to the order Rhizophydiales, that are host-specific chytrids that infect various phytoplankton species, mostly diatoms (Jobard, Rasconi, and Sime-Ngando 2010; Rasconi, Jobard, and Sime-Ngando 2011). The prevalence of Rhizophydiales in the Sanabria Lake ecosystem was not surprising given that they are the most common planktonic chytrids in lacustrine ecosystems (Monchy et al. 2011). A species of *Rhizophydiales* was probably the causative agent of a chytrid infection in Sanabria Lake in 2014 that diminished the population of the diatom *Tabellaria fenestrata* and controlled an algal bloom caused by eutrophication (Llorente and Seoane 2020). The relative abundance of Perkinsea, a group of parasitic alveolates, ranged from 13% to 18% of total parasite abundance across the different sampling sites. Perkinsea were previously considered as strictly marine parasites (Norén, Moestrup, and Rehnstam-Holm 1999; Erard-Le Denn, Chrétiennot-Dinet, and Probert 2000; Villalba et al. 2004; Figueroa et al. 2008; Leander and Hoppenrath 2008) until molecular environmental surveys revealed an unprecedented diversity of these organisms in freshwaters (Richards et al. 2005; Lefèvre et al. 2008; Lepère, Domaizon, and Debroas 2008; Bråte et al. 2010).

The parasitic community of each sampling site carried a unique taxonomic signature. The parasitic community of the tributary stream was characterised by a higher proportion of Apicomplexa (17%) and Labyrinthulomycetes (12%) in comparison to the other sampling sites. Most of the apicomplexan ASVs in the tributary stream fell into eugregarines, the most abundant apicomplexan group in environmental surveys (del Campo et al. 2019). Parasitic Stramenopiles (Pseudofungi), a significant component of freshwater ecosystems (Cooper, Pillinger, and Ridge 1997), constituted the second most abundant group in Laguna and represented 20% of the Stramenopiles and 7% (76,070 reads) of all eukaryotes in this small pond (Supplementary Figure 6). Within the group of parasitic Stramenopiles (Supplementary Figure 6), there was observed a higher prevalence of Oomycetes that are common fish pathogens (van West 2006; Phillips et al. 2008) in Laguna in comparison to the other sampling sites. Finally, Sanabria Lake harboured a higher relative abundance of Ichthyosporea (12%, 96,491 reads) in comparison to the other sampling sites (Stream: 2%, Laguna: 1%). The majority of the Ichthyosporea in Sanabria were associated with the marine genera *Abeoforma* (69%), *Sphaeroforma* (17%) and *Pseudoperkinsus* (10%), none of them previously identified in a freshwater environment.

To confirm the identity of the ichthyosporean ASVs in Sanabria Lake we analysed them by phylogenetic placement. We compiled a dataset that encompassed all the extant diversity of unicellular Holozoa (n=234). Half of the complete 18S rDNA gene sequences used to build the reference tree belonged to uncultured environmental taxa. A total of 132 ASVs identified as Ichthyosporea by the Ribosomal Database Project (RDP) classifier were placed into the 465 branches of the reference tree (Supplementary Figure 8). Most of the queries were placed in a clade formed by the freshwater anuran parasite *Anurofeca richardsi*, the marine *Creolimax fragrantissima, Pseudoperkinsus tapetis* and *Sphaeroforma arctica*, and some uncultured environmental taxa (Supplementary Figure 8). The 132 ichthyosporean queries were clustered into 17 clades in the best-hit placement tree (Supplementary Figure 9). Most of the clades were associated with freshwater sequences. Clade 4, the one formed by the larger number of sequences, was assigned to the FRESHIP2 group (del Campo and Ruiz-Trillo, 2013), expanding the known molecular diversity of these environmental taxa. Clades 13, 14 and 15 were assigned to *Anurofeca richardsi* and clade 9 to *Caullerya mensii*, another freshwater parasite that infects *Daphnia pulex* (Lu et al. 2020). We identified two clades that were directly associated with marine Ichthyosporea, clade 6 that branched as sister to *Abeoforma whisleri* and clade 16 that branched as sister to *Sphaeroforma arctica* (Supplementary Figure 9). The genera *Abeoforma* and *Sphaeroforma* were previously considered exclusively marine and this is the first record that connects these taxa to freshwater habitats. As freshwater habitats are increasingly explored by molecular means, the number of taxa that have been previously reported as exclusively marine and later were found in freshwater surveys continues to expand (Bråte et al. 2010; Simon et al. 2015; Richards and Bass 2005; Annenkova, Giner, and Logares 2020; Massana et al. 2006; Simon et al. 2015; Yi et al. 2017; Mukherjee et al. 2019).

### Abundant and potentially novel freshwater microbial eukaryotes

Metabarcoding biodiversity studies have shown that great part of the extant microbial diversity remains undocumented (Pawlowski et al. 2012; del Campo et al. 2014). In a metabarcoding survey, a species can be described as novel either because it is completely unknown to science or because the particular molecular marker database used in the study does not include available information on the species. In this study, we used the term molecular novelty to define any organism whose V4 hypervariable region of the 18S rRNA gene is not present in our reference database without discriminating between the two aforementioned cases.

To check whether Sanabria Lake and its surrounding freshwater systems could be a potential sampling site to isolate new organisms, we investigated the molecular novelty by first selecting potentially novel ASVs. We used the Ribosomal Database Project (RDP) classifier (Wang et al. 2007) to assign taxonomy to the ASVs. The RDP classifier provides for each ASV an assignment of the best matching taxa together with a bootstrap confidence score at each taxonomic rank. This score represents the level of confidence of the taxonomic assignment at each rank, from supergroup to species. Here, we define as poorly assigned, thus potentially novel, all ASVs with bootstrap confidence score value <97 at the supergroup level. We were interested in identifying the most abundant and novel microbial eukaryotes in our study site, so we selected all ASVs with more than 1,000 reads and bootstrap confidence score value lower than the aforementioned established novelty threshold.

To assign taxonomy to the queries of our dataset, we analysed them using phylogenetic placement (Figure 5). We first constructed a comprehensive eukaryotic reference tree with 618 eukaryotic taxa that encompassed all the extant eukaryotic diversity according to the latest classification of eukaryotes (Adl et al. 2019). We designed the reference tree with two criteria. First, to be inclusive in order to minimise phylogenetic placement artefacts related to taxonomic sampling and second to be non-redundant in order to be smaller and thus easier to handle in the post placement analyses. The amplicon short sequences were aligned to the reference alignment and the amplicon sequences that were not aligned in the V4 region were removed as artefacts after manual inspection. We placed a total of 113 ASV V4 queries into 1,233 branches of the reference tree.

**Figure 5.**
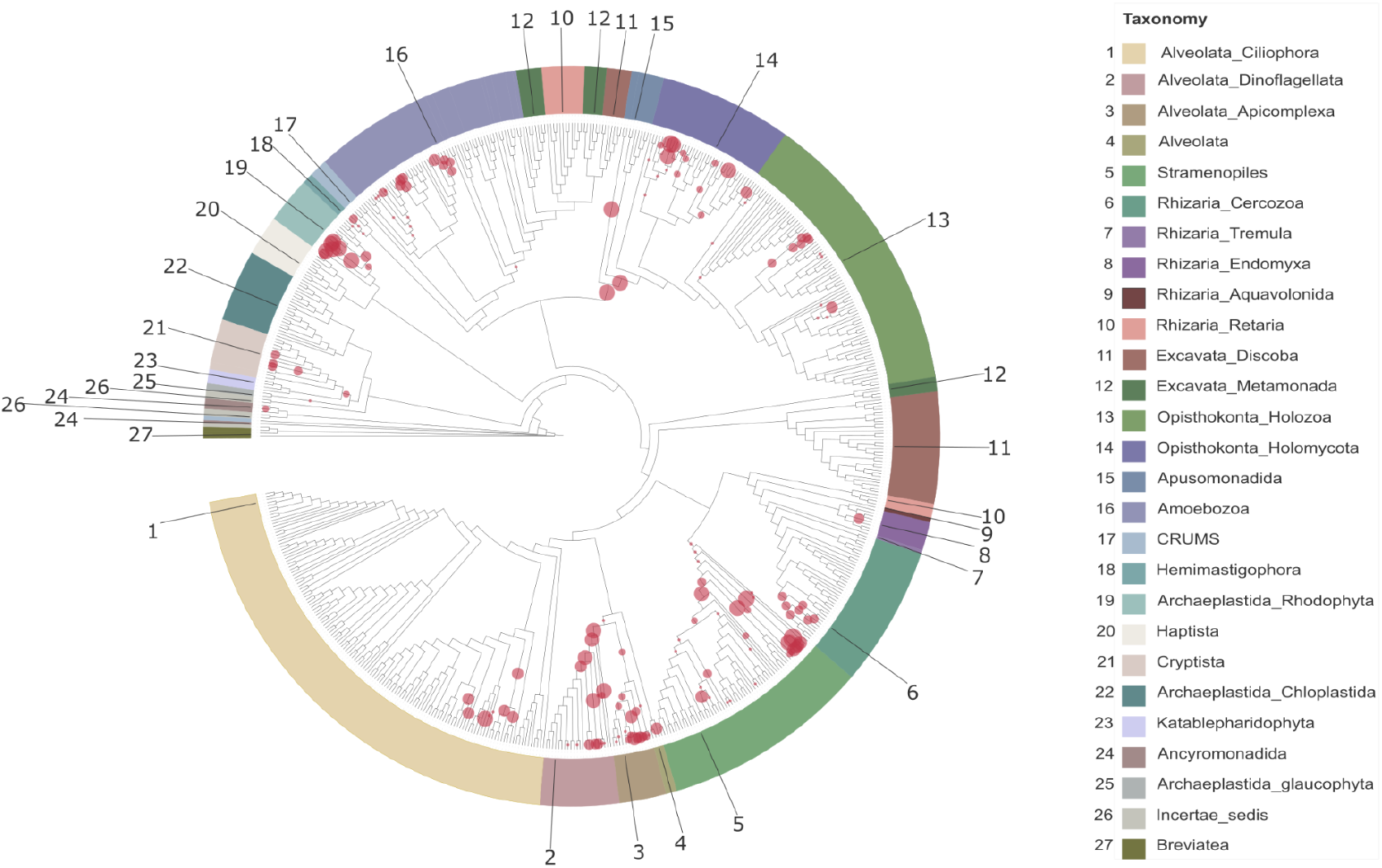
Phylogenetic placement of 113 ASVs into 1233 branches of a reference tree with 618 taxa that span all extant eukaryotic diversity as described in Adl et al. 2019. The ASVs have bootstrap confidence score value <97 at the supergroup level in the RDP classifier (see Methods) and total abundance greater than 1000 reads. The diameter of the circles indicates the number of ASVs placed in the branch. One ASV can be placed on multiple branches until it reaches accumulated likelihood_weight_ratio 1.

Most of the ASV placements in the tree were found in the leaf nodes of Rhodophyta (Archaeplastida), Bigyra (Stramenopiles), early-branching Nucletmycea (also known as Holomycota) and Apicomplexa (Alveolata) pinpointing these clades as parts of the tree with potential novel undescribed molecular diversity (Figure 5). An elevated number of placements was spotted in the internal nodes of Dinophyta and the divergence between Opisthokonta and Apusomonadida (Figure 5). Apusomonadida is a recently defined phylum with a key phylogenetic position to understand the origin of the eukaryotic cell. Apusomonads are rarely detected in environmental studies (Purificación López-García et al. 2003; Not et al. 2008; Takishita et al. 2007; W. Orsi et al. 2012; Lesaulnier et al. 2008) and can be considerably more diverse than is currently perceived (Torruella, Moreira, and López-García 2017). We report previously undocumented diversity associated with the genera *Cryptomonas* and *Chilomonas* inside Cryptista, the naked filose amoebae of the genus *Vampyrella* (Endomyxa) and the frequently detected by 18S rRNA gene sequencing eukaryovorous biflagellate Aquavolon (Bass et al. 2018). No placement was recorded inside Excavata.

## Conclusions

Metabarcoding analyses of the V4 hypervariable region of the 18S rRNA gene from diverse habitats in Sanabria Lake and the surrounding freshwater ecosystems uncovered a rich and diverse microeukaryotic community. One fifth of the diversity of microeukaryotes identified in Sanabria Lake are parasites, stressing the importance of parasitic taxa in the freshwater ecosystems. Our observations regarding the taxonomic composition of the microeukaryotic community overlap with microscopical data based on morphology but expand the total biodiversity recorded in the lake by adding taxa that were either insufficiently abundant to be detected by traditional methods or inconspicuous due to lack of taxonomically informative morphological characters. Tributary stream samples were significantly more species-rich than samples from Sanabria lake and Laguna. We found that sediments harbour the greatest diversity among different habitats. We observed compositional heterogeneity among the microbial communities of Sanabria Lake and the surrounding freshwater ecosystem. Phylogenetic placement analysis showed that Sanabria Lake and the surrounding freshwater ecosystems would be good targets for future studies aiming the discovery of potential novel microeukaryotes. This is the first metabarcoding record of the diversity in Sanabria Lake. Our results expand our understanding of the microbial communities in oligomesotrophic, temperate, lacustrine ecosystems and can be used as reference for future studies in the area.

## Acknowledgments

This work was supported by grants (BFU2017-90114-P and PID2020-120609GB-I00) funded by MCIN/AEI/ 10.13039/501100011033 and “ERDF A way of making Europe”. It has also received funding from the European Union’s Horizon 2020 research and innovation programme under the Marie Skłodowska-Curie grant agreement no. H2020-MSCA-ITN-2015-675752 (SINGEK Project, http://www.singek.eu/). This project has also received funding from the European Research Council (ERC) under the European Union’s Horizon 2020 research and innovation programme (grant agreement No. 949745) and the support of a fellowship from “la Caixa” Foundation (ID 100010434) with fellowship code LCF/BQ/PI19/11690008.

## Data accessibility

Raw data are available in the European Nucleotide Archive (ENA) under the accession number PRJEB23911.

## Authors’ contributions

K.M. designed and conducted the data analyses, interpreted the results, designed the figures and wrote the draft and the revised manuscript. A.S.A., D.L.E, M.A., and A.G. collected the data and revised the manuscript. D.J.R. interpreted the results and revised the manuscript. I.R.-T. conceived and designed the study, supervised the work and revised the manuscript. All authors approved the final version of the manuscript and agreed to be accountable for all aspects of the work in ensuring that questions related to the accuracy or integrity of any part of the work are appropriately investigated and resolved.

