## Supplementary Material for "Taxonomic composition, community structure and molecular novelty of microeukaryotes in a temperate oligomesotrophic lake as revealed by metabarcoding"

### Supplementary Figures

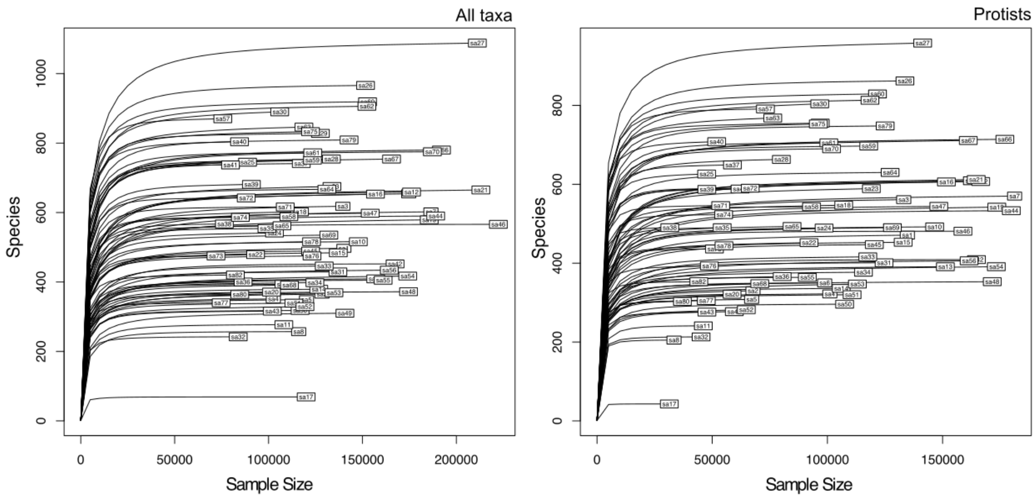


**Supplementary Figure 1.** Rarefaction curves for the 81 samples of the study. Rarefaction curves were drawn for all taxa (dataset D2) and for protists (dataset D3). For a detailed description of the datasets see Supplementary Table 3.

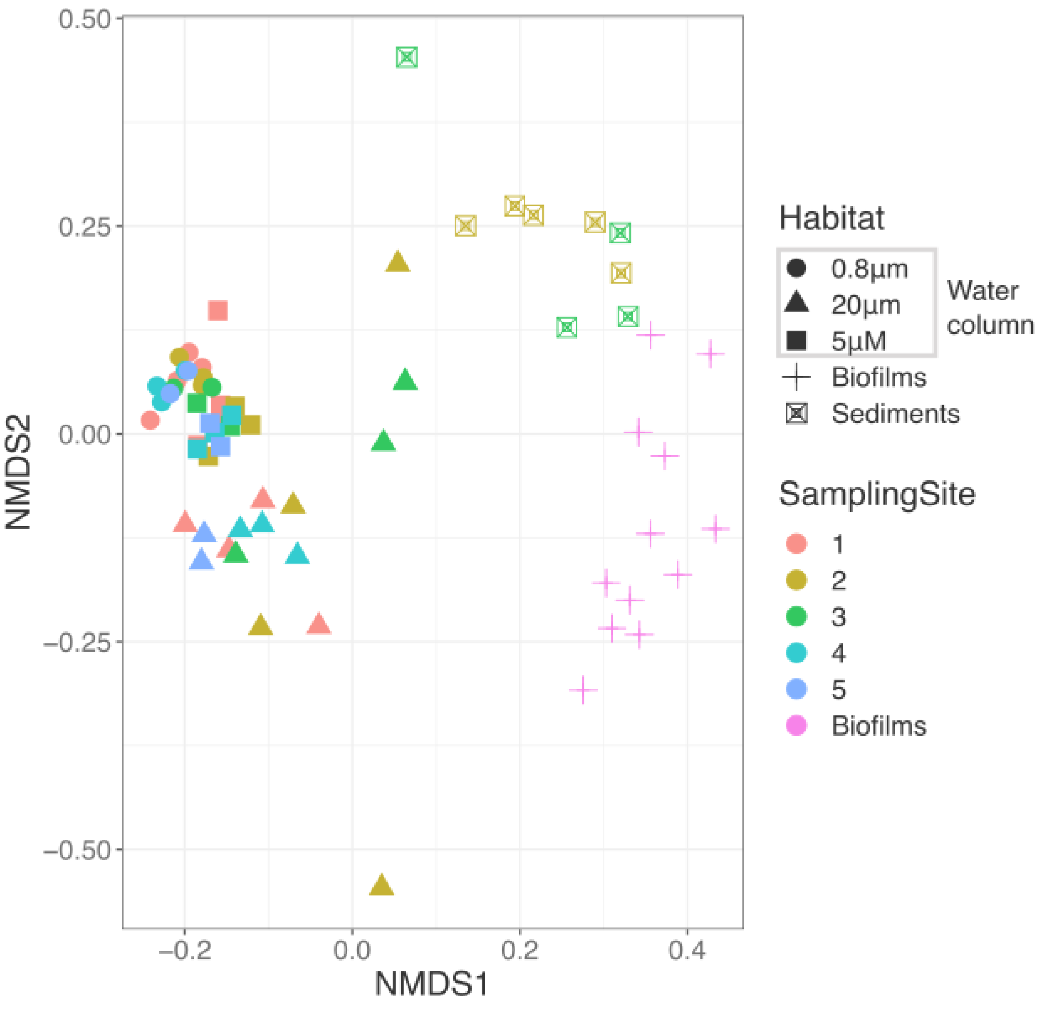


**Supplementary Figure 2.** Reduced-space NMDS plot showing microbial eukaryotes community structure based on Bray–Curtis dissimilarity. The dissimilarity matrix was calculated from the rarefied at even depth (31361 reads) abundances of protist ASVs present only in Sanabria samples (dataset D5) (Stress = 0.1679787, Procrustes: rmse 0.001014908 max resid 0.006658075)

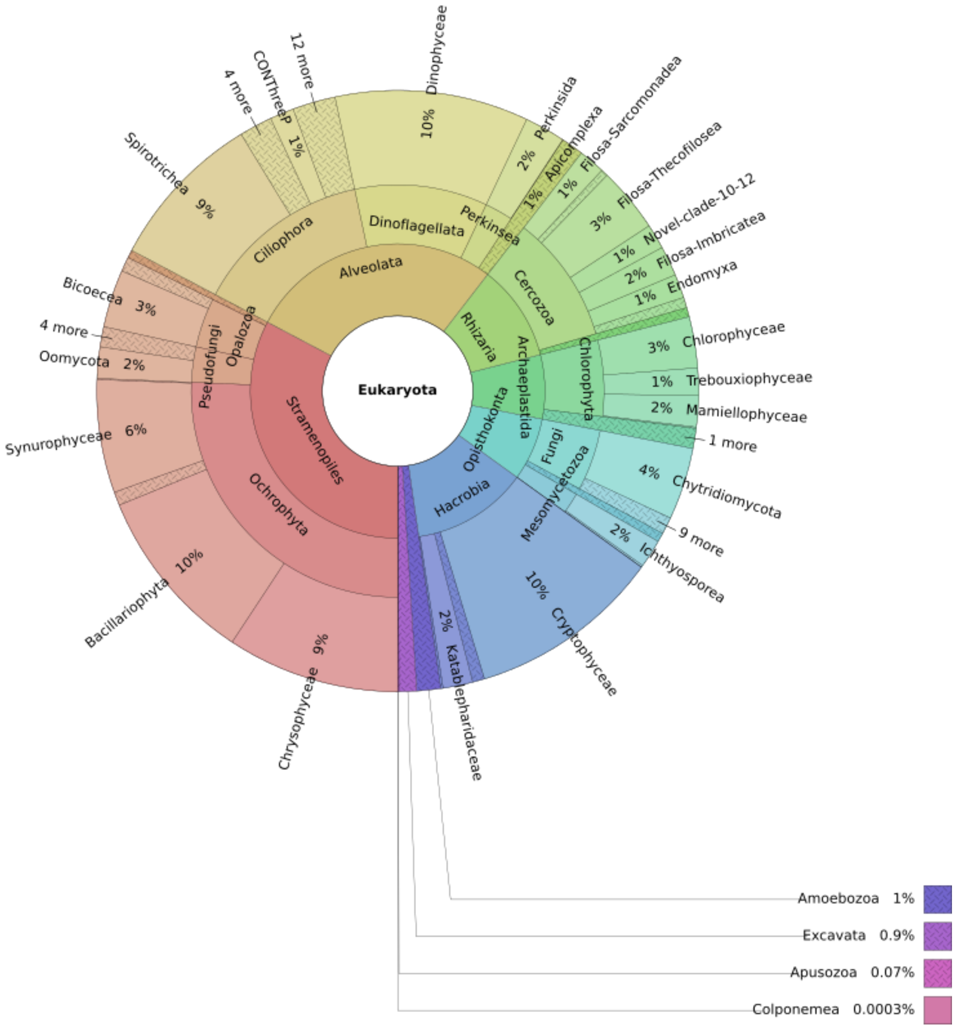


**Supplementary Figure 3.** KRONA representation of the eukaryotic microbial diversity as inferred by DADA2 at division level in Sanabria Lake (samples S1-S5). (Total reads: 6375317)

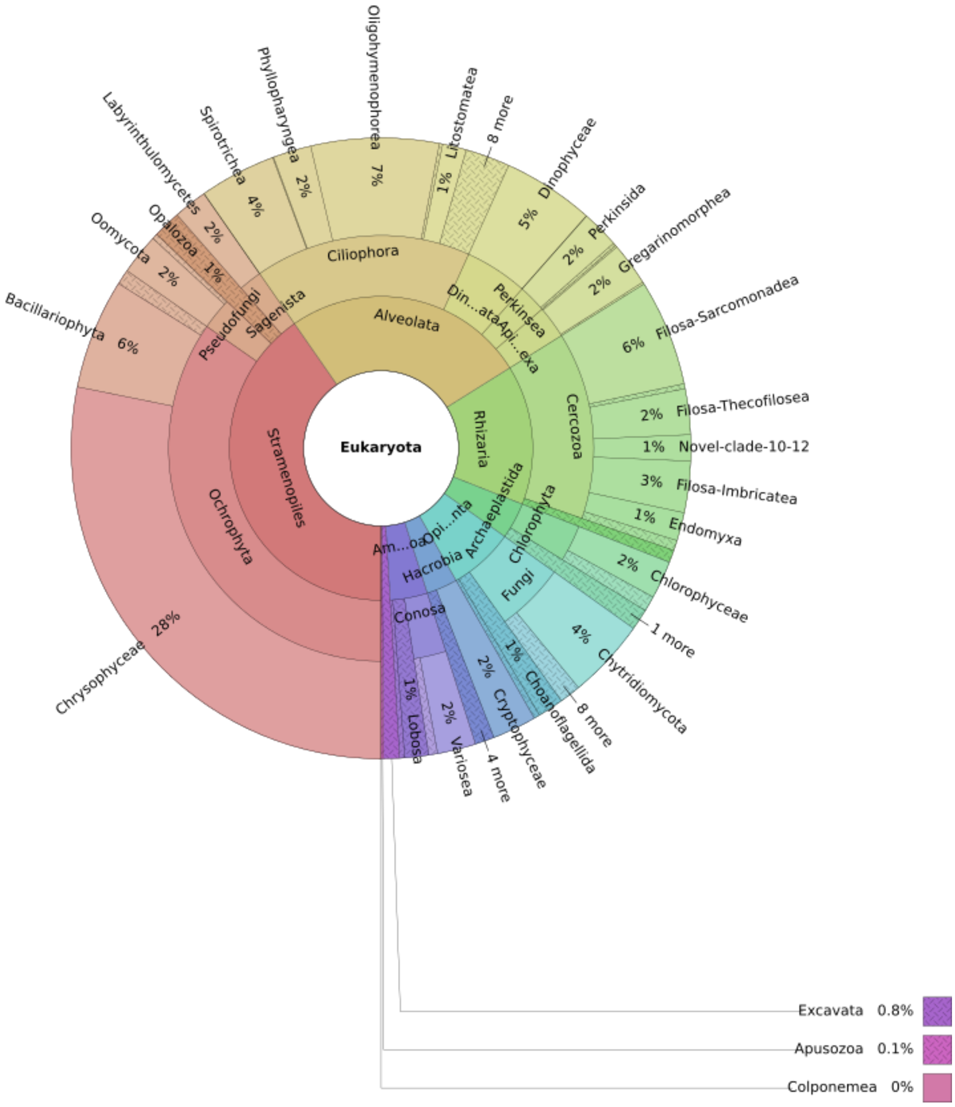


**Supplementary Figure 4.** KRONA representation of the eukaryotic microbial diversity as inferred by DADA2 at division level in the tributary stream (samples S6-S7). (Total reads: 701233)

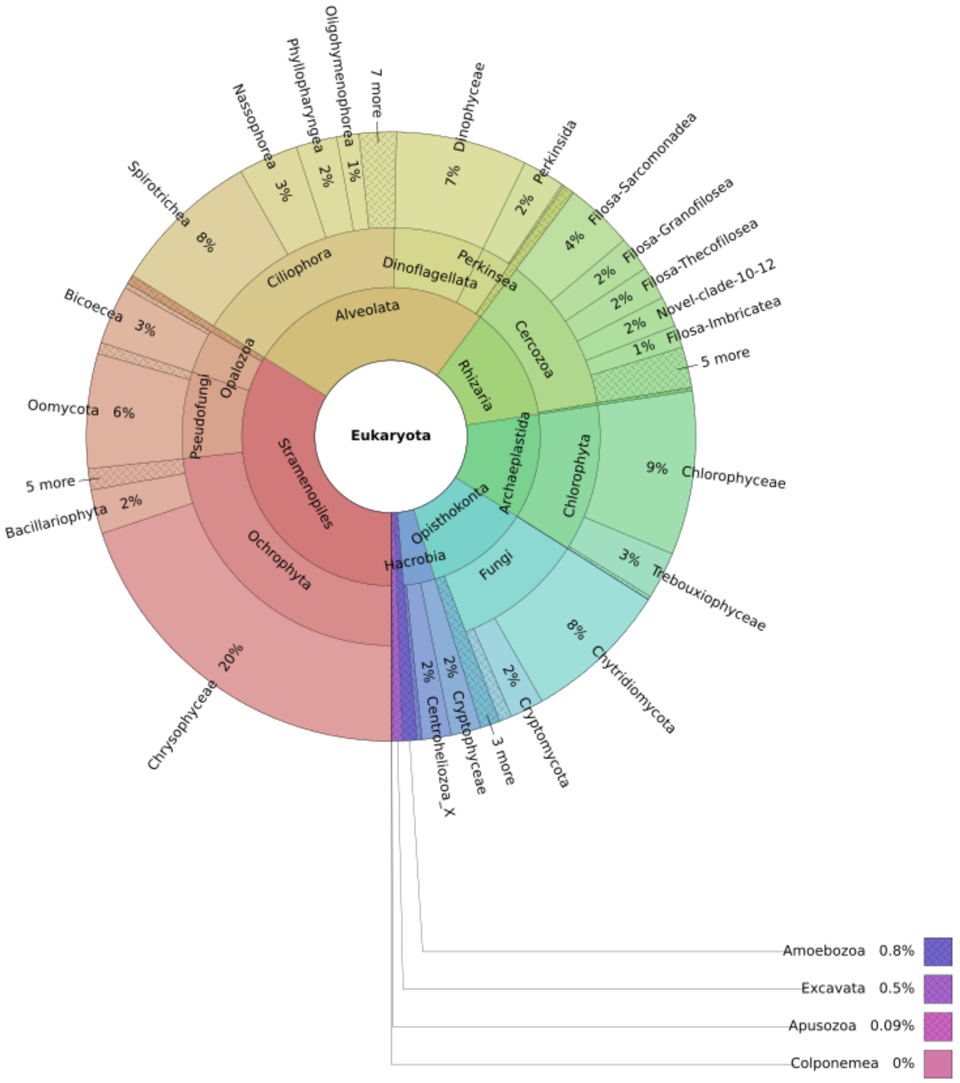


**Supplementary Figure 5.** KRONA representation of the eukaryotic microbial diversity as inferred by DADA2 at division level in the nearby pond (Laguna) (samples S8-S10). (Total reads: 1138974)

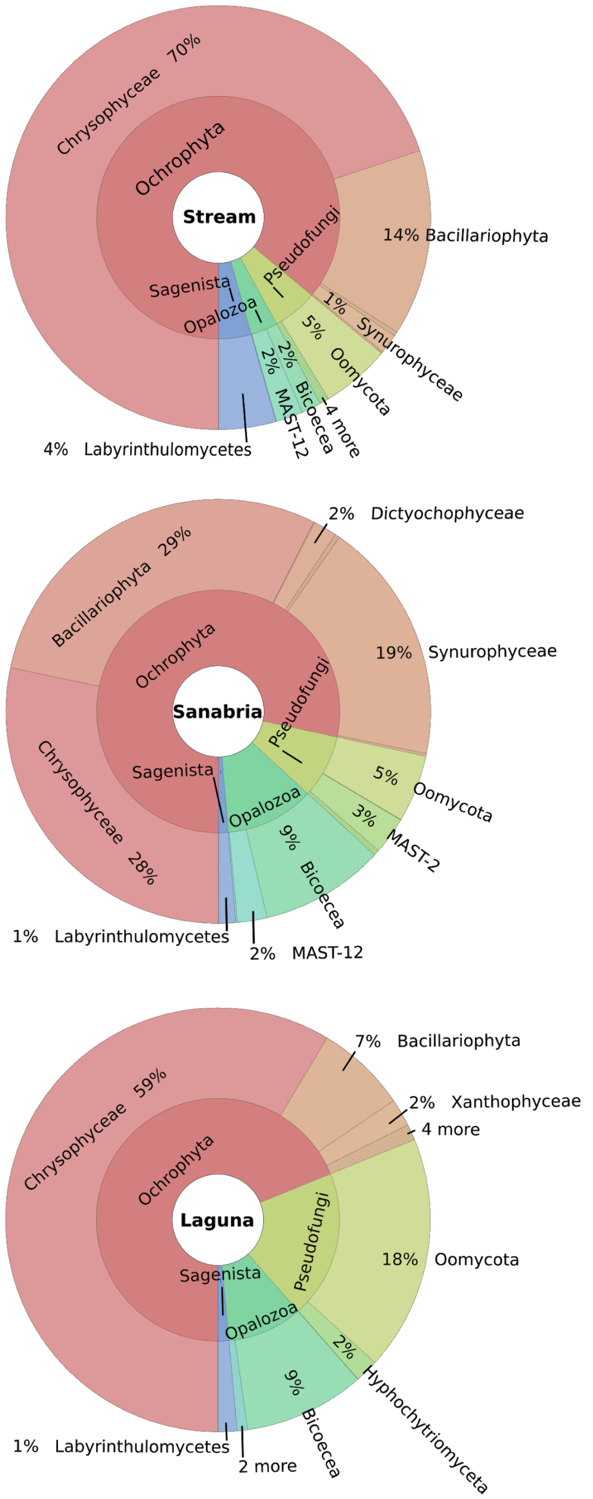


**Supplementary Figure 6.** KRONA representation of the Stramenopiles diversity as inferred by ASV in the three sampling sites.

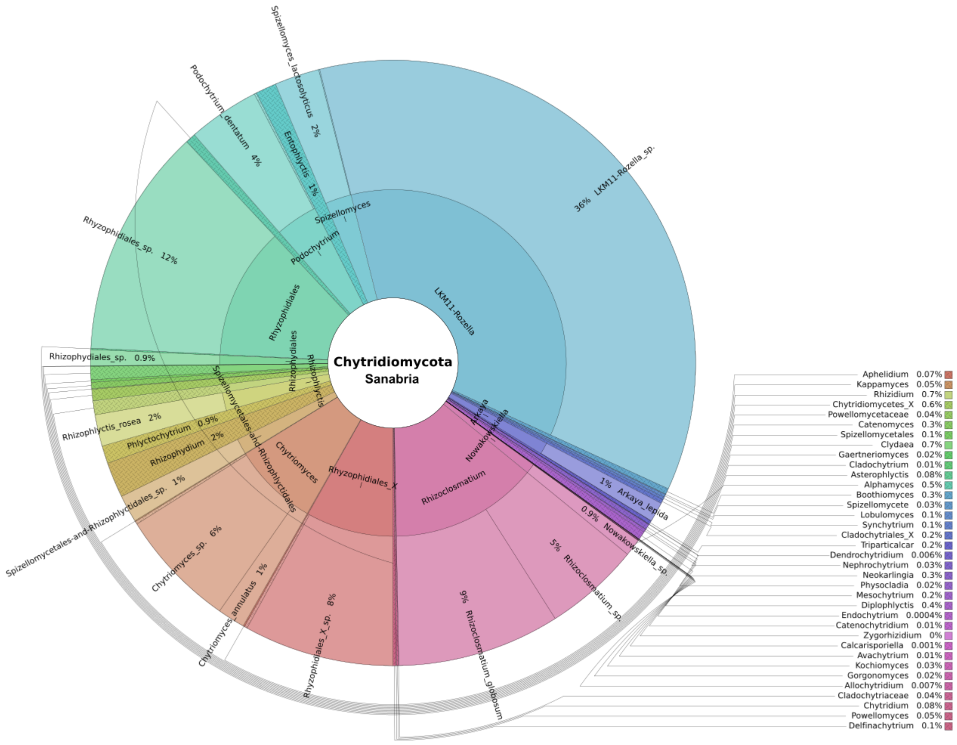


**Supplementary Figure 7.** KRONA representation of the Chytridiomycota diversity as inferred by ASV in the Sanabria Lake. (Total reads: 243311)

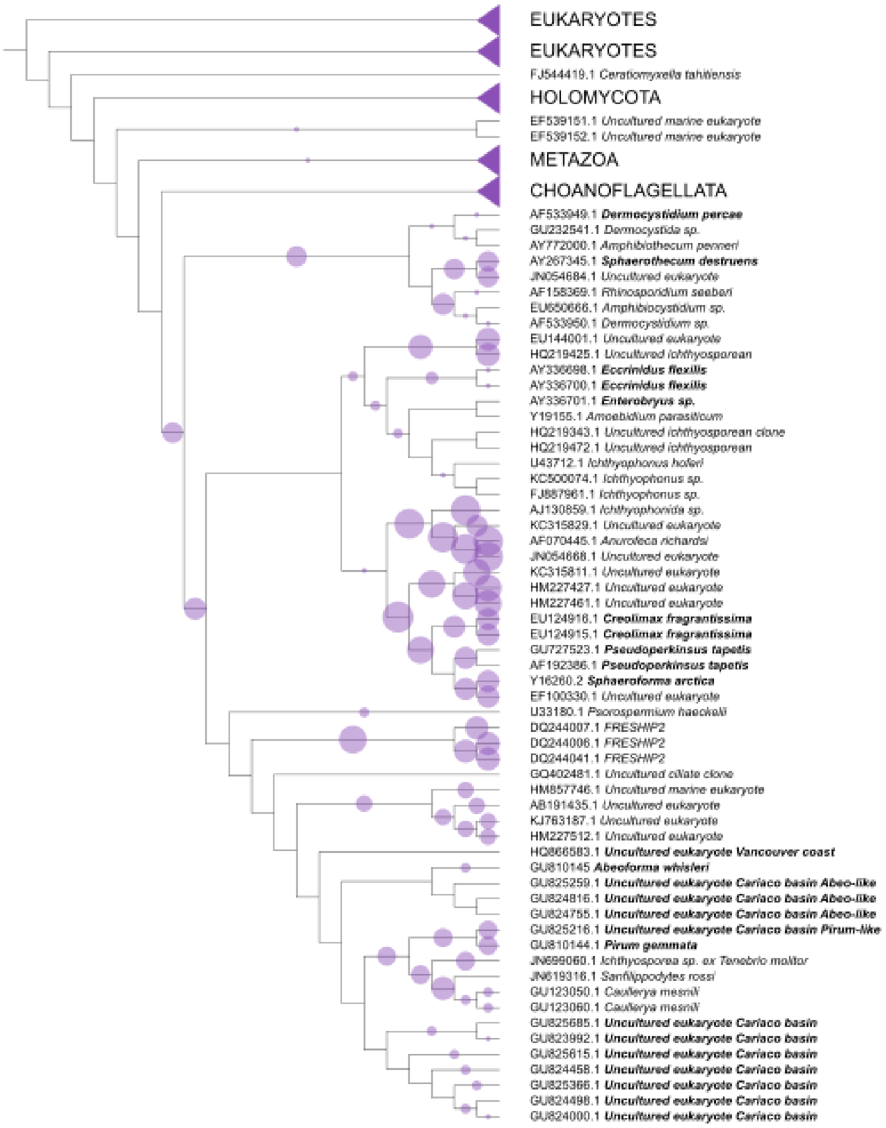


**Supplementary Figure 8.** Phylogenetic placement of 132 ASVs identified as Ichthyosporea into 465 branches of a reference tree with 234 taxa. Marine taxa are bold. Nodes that do not include any placements were collapsed. The diameter of the circles indicates the number of ASVs placed in the branch. One ASV can be placed to multiple branches until it reaches accumulated like_weight_ratio 1.

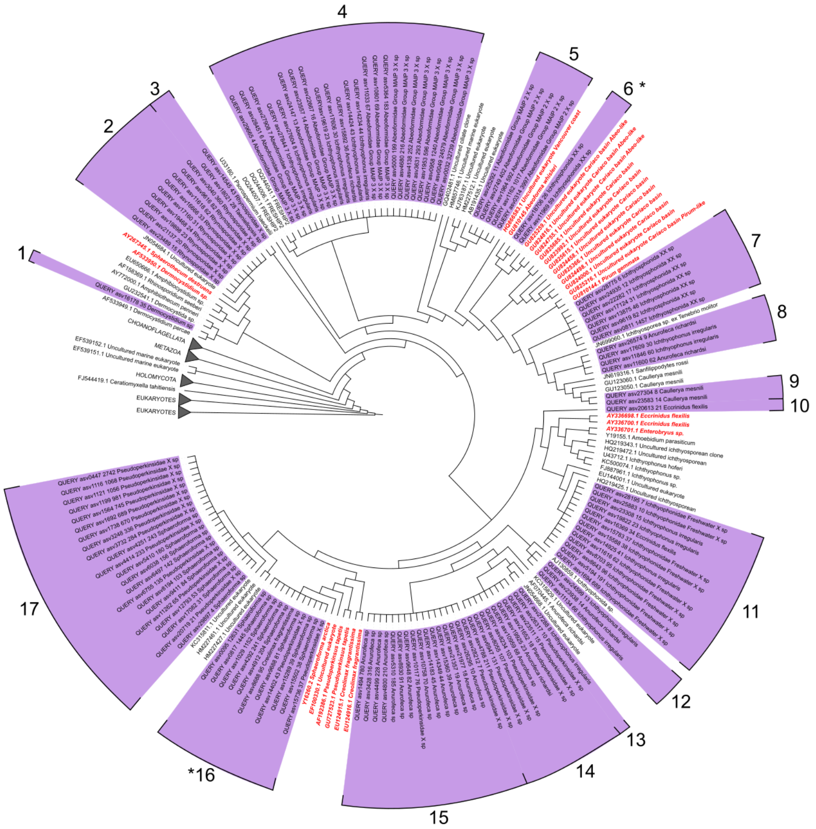


**Supplementary Figure 9.** Best-hit placement tree of 132 Ichthyosporea ASVs into 465 branches of a reference tree with 234 taxa that encompass the extant holozoan diversity with emphasis on environmental uncultured taxa. Queries are marked purple. The queries name includes read abundance and the taxonomy assigned by the RDP classifier. Reference taxa are black and reference marine taxa are marked red. Each query is uniquely placed in the position with the highest likelihood–weight ratio. Numbers represent the clades of clustered ASVs and asterisks denote marine associated clades.

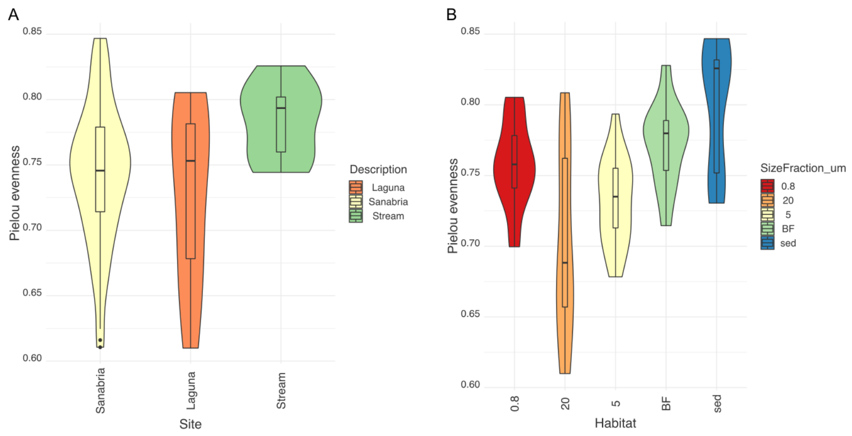


**Supplementary Figure 10.** Pielou's evenness across (A) the sampling sites (A) and (B) the different habitats of Sanabria Lake (excluding the surrounding freshwater habitats) (B) as calculated by the ASVs rarefied at 31361 reads .

### Supplementary Tables

| Sample | Site | Sampling Site | SamplingDepth (m) | Depth_Range | Chl_max | SizeFraction_µm | Hour | Habitat | Tempera-ture (°C) |
|---|---|---|---|---|---|---|---|---|---|
| sa1 | Sanabria | 1 | 14 | deep | no | 0.8 | morning | wc | 6.6 |
| sa2 | Sanabria | 1 | 14 | deep | no | 20 | morning | wc | 6.6 |
| sa3 | Sanabria | 1 | 14 | deep | no | 5 | morning | wc | 6.6 |
| sa4 | Sanabria | 1 | 35 | deep | no | 0.8 | morning | wc | 6.2 |
| sa5 | Sanabria | 1 | 35 | deep | no | 20 | morning | wc | 6.2 |
| sa6 | Sanabria | 1 | 35 | deep | no | 5 | morning | wc | 6.2 |
| sa7 | Sanabria | 1 | 4 | shallow | yes | 0.8 | morning | wc | 7.5 |
| sa8 | Sanabria | 1 | 4 | shallow | yes | 20 | morning | wc | 7.5 |
| sa9 | Sanabria | 1 | 4 | shallow | yes | 5 | morning | wc | 7.5 |
| sa10 | Sanabria | 1 | sur | sur | no | 0.8 | morning | wc | 7.6 |
| sa11 | Sanabria | 1 | sur | sur | no | 20 | morning | wc | 7.6 |
| sa12 | Sanabria | 1 | sur | sur | no | 5 | morning | wc | 7.6 |
| sa13 | Laguna | 10 | NA | NA | no | 0.8 | morning | wc | NA |
| sa14 | Laguna | 10 | NA | NA | no | 20 | morning | wc | NA |
| sa15 | Laguna | 10 | NA | NA | no | 5 | morning | wc | NA |
| sa16 | Sanabria | 2 | 12 | deep | no | 0.8 | morning | wc | 6.5 |
| sa17 | Sanabria | 2 | 12 | deep | no | 20 | morning | wc | 6.5 |
| sa18 | Sanabria | 2 | 12 | deep | no | 5 | morning | wc | 6.5 |
| sa19 | Sanabria | 2 | 2 | shallow | yes | 0.8 | morning | wc | 6.9 |
| sa20 | Sanabria | 2 | 2 | shallow | yes | 20 | morning | wc | 6.9 |
| sa21 | Sanabria | 2 | 2 | shallow | yes | 5 | morning | wc | 6.9 |
| sa22 | Sanabria | 2 | sur | sur | no | 0.8 | morning | wc | 7.1 |
| sa23 | Sanabria | 2 | sur | sur | no | 20 | morning | wc | 7.1 |
| sa24 | Sanabria | 2 | sur | sur | no | 5 | morning | wc | 7.1 |
| sa25 | Sanabria | 2 | sed | sed | no | sed | morning | sed | 6.2 |
| sa26 | Sanabria | 2 | sed | sed | no | sed | morning | sed | 6.2 |
| sa27 | Sanabria | 2 | sed | sed | no | sed | morning | sed | 6.2 |
| sa28 | Sanabria | 2 | sed | sed | no | sed | morning | sed | 6.2 |
| sa29 | Sanabria | 2 | sed | sed | no | sed | morning | sed | 6.2 |
| sa30 | Sanabria | 2 | sed | sed | no | 20 | morning | sed | 6.2 |
| sa31 | Sanabria | 3 | 1 | shallow | yes | 0.8 | noon | wc | 7.9 |
| sa32 | Sanabria | 3 | 1 | shallow | yes | 20 | noon | wc | 7.9 |
| sa33 | Sanabria | 3 | 1 | shallow | yes | 5 | noon | wc | 7.9 |
| sa34 | Sanabria | 3 | 5 | shallow | no | 0.8 | noon | wc | 7.0 |
| sa35 | Sanabria | 3 | 5 | shallow | no | 20 | noon | wc | 7.0 |
| sa36 | Sanabria | 3 | 5 | shallow | no | 5 | noon | wc | 7.0 |
| sa37 | Sanabria | 3 | sed | sed | no | 20 | noon | sed | 6.9 |
| sa38 | Sanabria | 3 | sed | sed | no | sed | noon | sed | 6.9 |
| sa39 | Sanabria | 3 | sed | sed | no | sed | noon | sed | 6.9 |
| sa40 | Sanabria | 3 | sed | sed | no | sed | noon | sed | 6.9 |

| | | | | | | | | | |
|---|---|---|---|---|---|---|---|---|---|
| sa41 | Sanabria | 3 | sed | sed | no | sed | noon | sed | 6.9 |
| sa42 | Sanabria | 4 | 25 | deep | no | 0.8 | afternoon | wc | 6.1 |
| sa43 | Sanabria | 4 | 25 | deep | no | 20 | afternoon | wc | 6.1 |
| sa44 | Sanabria | 4 | 25 | deep | no | 5 | afternoon | wc | 6.1 |
| sa45 | Sanabria | 4 | 9 | shallow | yes | 0.8 | afternoon | wc | 7.0 |
| sa46 | Sanabria | 4 | 9 | shallow | yes | 20 | afternoon | wc | 7.0 |
| sa47 | Sanabria | 4 | 9 | shallow | yes | 5 | afternoon | wc | 7.0 |
| sa48 | Sanabria | 4 | sur | sur | no | 0.8 | afternoon | wc | 8.3 |
| sa49 | Sanabria | 4 | sur | sur | no | 20 | afternoon | wc | 8.3 |
| sa50 | Sanabria | 4 | sur | sur | no | 5 | afternoon | wc | 8.3 |
| sa51 | Sanabria | 5 | 10 | deep | yes | 0.8 | afternoon | wc | 8.1 |
| sa52 | Sanabria | 5 | 10 | deep | yes | 20 | afternoon | wc | 8.1 |
| sa53 | Sanabria | 5 | 10 | deep | yes | 5 | afternoon | wc | 8.1 |
| sa54 | Sanabria | 5 | sur | sur | no | 0.8 | afternoon | wc | 8.4 |
| sa55 | Sanabria | 5 | sur | sur | no | 20 | afternoon | wc | 8.4 |
| sa56 | Sanabria | 5 | sur | sur | no | 5 | afternoon | wc | 8.4 |
| sa57 | Stream | 6 | sur | sur | no | 0.8 | morning | wc | NA |
| sa58 | Stream | 6 | sur | sur | no | 20 | morning | wc | NA |
| sa59 | Stream | 6 | sur | sur | no | 5 | morning | wc | NA |
| sa60 | Stream | 7 | sed | sed | no | sed | morning | sed | NA |
| sa61 | Stream | 7 | sed | sed | no | sed | morning | sed | NA |
| sa62 | Stream | 7 | sed | sed | no | sed | morning | sed | NA |
| sa63 | Stream | 7 | sed | sed | no | sed | morning | sed | NA |
| sa64 | Laguna | 8 | NA | NA | no | 0.8 | morning | wc | NA |
| sa65 | Laguna | 8 | NA | NA | no | 20 | morning | wc | NA |
| sa66 | Laguna | 8 | NA | NA | no | 5 | morning | wc | NA |
| sa67 | Laguna | 9 | NA | NA | no | 0.8 | morning | wc | NA |
| sa68 | Laguna | 9 | NA | NA | no | 20 | morning | wc | NA |
| sa69 | Laguna | 9 | NA | NA | no | 5 | morning | wc | NA |
| sa70 | Sanabria | bf | bf | bf | no | bf | noon | bf | NA |
| sa71 | Sanabria | bf | bf | bf | no | bf | noon | bf | NA |
| sa72 | Sanabria | bf | bf | bf | no | bf | noon | bf | NA |
| sa73 | Sanabria | bf | bf | bf | no | bf | noon | bf | NA |
| sa74 | Sanabria | bf | bf | bf | no | bf | noon | bf | NA |
| sa75 | Sanabria | bf | bf | bf | no | bf | noon | bf | NA |
| sa76 | Sanabria | bf | bf | bf | no | bf | noon | bf | NA |
| sa77 | Sanabria | bf | bf | bf | no | bf | noon | bf | NA |
| sa78 | Sanabria | bf | bf | bf | no | bf | noon | bf | NA |
| sa79 | Sanabria | bf | bf | bf | no | bf | noon | bf | NA |
| sa80 | Sanabria | bf | bf | bf | no | bf | noon | bf | NA |
| sa81 | Sanabria | bf | bf | bf | no | bf | noon | bf | NA |
| sa82 | Sanabria | bf | bf | bf | no | bf | noon | bf | NA |

**Supplementary Table 1** Sampling metadata. The Sampling Site variable refers to the sites marked with orange triangles in Figure 1. Chl_max denotes whether the sample was collected in the chlorophyll maximum point or not. Size_fraction_μm variable refers to the filter size used to filter each sample (when applicable). Abbreviations : bf = biofilm, sed = sediment, wc = water column, sur = surface

| sample | input | filtered | denoisedF | denoisedR | merged | nonchim |
|---|---|---|---|---|---|---|
| sa1 | 233721 | 189443 | 174882 | 183756 | 6298 | 6111 |
| sa2 | 225677 | 178581 | 159411 | 175513 | 995 | 994 |
| sa3 | 237977 | 192981 | 177778 | 185814 | 9154 | 8491 |
| sa4 | 180795 | 145523 | 132286 | 137789 | 9622 | 9457 |
| sa5 | 197831 | 161002 | 139850 | 154955 | 2624 | 2622 |
| sa6 | 177518 | 146162 | 133484 | 139305 | 6966 | 6917 |
| sa7 | 299485 | 248210 | 233456 | 242129 | 5678 | 5444 |
| sa8 | 224661 | 179221 | 155080 | 176497 | 1323 | 1285 |
| sa9 | 273776 | 227988 | 214380 | 224301 | 3281 | 3266 |
| sa10 | 256256 | 208112 | 192418 | 200919 | 6279 | 6080 |
| sa11 | 241304 | 191051 | 169484 | 187815 | 747 | 624 |
| sa12 | 290441 | 236865 | 216577 | 229743 | 6599 | 6588 |
| sa13 | 256678 | 209165 | 199337 | 204955 | 15401 | 14976 |
| sa14 | 218771 | 182574 | 169251 | 179139 | 2382 | 2382 |
| sa15 | 225041 | 186609 | 174869 | 180912 | 7514 | 7327 |
| sa16 | 272938 | 223837 | 201307 | 212997 | 9751 | 9533 |
| sa17 | 235654 | 192947 | 163134 | 192342 | 781 | 781 |
| sa18 | 211074 | 170276 | 154404 | 162065 | 8225 | 8142 |
| sa19 | 322885 | 263074 | 242439 | 253955 | 7314 | 7101 |
| sa20 | 171946 | 141743 | 129551 | 139564 | 1020 | 1020 |
| sa21 | 377763 | 308603 | 280780 | 300926 | 7921 | 7920 |
| sa22 | 165000 | 136403 | 120840 | 127983 | 6545 | 6382 |
| sa23 | 217072 | 182855 | 166330 | 176715 | 3943 | 3943 |
| sa24 | 178373 | 148938 | 134664 | 142602 | 5682 | 5679 |
| sa25 | 239395 | 181125 | 127705 | 140860 | 15416 | 14886 |
| sa26 | 325594 | 253447 | 208569 | 223554 | 6332 | 6276 |
| sa27 | 453233 | 368265 | 314546 | 338099 | 9810 | 9626 |
| sa28 | 288153 | 228759 | 181464 | 197935 | 6510 | 6248 |
| sa29 | 218156 | 177599 | 158780 | 164295 | 3329 | 3329 |
| sa30 | 239023 | 186333 | 143991 | 155488 | 6420 | 6419 |
| sa31 | 227545 | 184268 | 171037 | 178608 | 3929 | 3929 |
| sa32 | 160733 | 129678 | 116318 | 127196 | 8666 | 8234 |
| sa33 | 232514 | 189788 | 176156 | 184474 | 4338 | 4254 |
| sa34 | 211193 | 172041 | 160116 | 166778 | 3748 | 3748 |
| sa35 | 173613 | 141075 | 120678 | 133211 | 1698 | 1698 |
| sa36 | 141502 | 114109 | 105843 | 111568 | 2613 | 2605 |
| sa37 | 199939 | 165233 | 138207 | 153542 | 2930 | 2878 |
| sa38 | 177541 | 138665 | 96418 | 110772 | 4838 | 4838 |
| sa39 | 206488 | 159979 | 118337 | 130392 | 9883 | 9390 |
| sa40 | 180967 | 144934 | 120037 | 132885 | 2006 | 2006 |
| sa41 | 179755 | 137100 | 102620 | 108761 | 12321 | 11857 |
| sa42 | 270646 | 222270 | 203513 | 213043 | 12474 | 12125 |
| sa43 | 175651 | 140943 | 124347 | 135978 | 2035 | 1859 |
| sa44 | 303280 | 250143 | 227395 | 238572 | 14219 | 13068 |
| sa45 | 216592 | 172117 | 155063 | 162552 | 6846 | 6648 |

| | | | | | | |
|---|---|---|---|---|---|---|
| sa46 | 332994 | 273957 | 250012 | 268713 | 21188 | 21028 |
| sa47 | 260304 | 209554 | 193625 | 202323 | 10590 | 10589 |
| sa48 | 288573 | 235347 | 226935 | 232721 | 4781 | 4572 |
| sa49 | 241141 | 198364 | 175188 | 193765 | 1379 | 1379 |
| sa50 | 196243 | 161225 | 150201 | 156643 | 3649 | 3649 |
| sa51 | 194835 | 158163 | 148110 | 153084 | 4914 | 4805 |
| sa52 | 193789 | 159382 | 144974 | 156297 | 2018 | 1925 |
| sa53 | 219487 | 180943 | 166597 | 175907 | 3632 | 3632 |
| sa54 | 291411 | 237462 | 223966 | 230724 | 6328 | 6117 |
| sa55 | 264376 | 220133 | 196525 | 215702 | 1932 | 1932 |
| sa56 | 272665 | 224623 | 208214 | 218602 | 12779 | 12717 |
| sa57 | 151803 | 123128 | 102038 | 111518 | 6719 | 6619 |
| sa58 | 169953 | 142715 | 134393 | 137734 | 8008 | 7271 |
| sa59 | 196094 | 165025 | 154967 | 160890 | 8729 | 7801 |
| sa60 | 299140 | 242938 | 207241 | 223298 | 3725 | 3421 |
| sa61 | 243050 | 193497 | 162462 | 173461 | 3545 | 3173 |
| sa62 | 303703 | 239176 | 193734 | 207300 | 3602 | 3424 |
| sa63 | 253357 | 200283 | 163001 | 178641 | 2566 | 2494 |
| sa64 | 207846 | 169867 | 158020 | 162857 | 2796 | 2712 |
| sa65 | 184216 | 150893 | 138926 | 144333 | 2860 | 2835 |
| sa66 | 338723 | 273929 | 256081 | 265254 | 13766 | 13703 |
| sa67 | 290991 | 240786 | 217243 | 230497 | 4011 | 4009 |
| sa68 | 225315 | 187749 | 167143 | 183108 | 2635 | 2598 |
| sa69 | 224752 | 184427 | 168697 | 179578 | 11333 | 11305 |
| sa70 | 299901 | 248549 | 228256 | 239169 | 2506 | 2506 |
| sa71 | 188868 | 153672 | 139122 | 147095 | 5365 | 5089 |
| sa72 | 155354 | 122784 | 108812 | 114998 | 2462 | 2361 |
| sa73 | 144906 | 116307 | 92574 | 100969 | 1255 | 1255 |
| sa74 | 164581 | 133580 | 113620 | 122205 | 1224 | 1224 |
| sa75 | 230701 | 190043 | 160740 | 172072 | 2359 | 2358 |
| sa76 | 204897 | 173716 | 161503 | 169392 | 4795 | 3823 |
| sa77 | 130698 | 102977 | 94986 | 99924 | 1673 | 1651 |
| sa78 | 229977 | 191065 | 174560 | 186214 | 3078 | 3013 |
| sa79 | 278817 | 218132 | 201467 | 209964 | 4520 | 4442 |
| sa80 | 152122 | 121619 | 110954 | 118101 | 1595 | 1595 |
| sa81 | 2248 | 1519 | 671 | 1275 | 0 | 0 |
| sa82 | 167108 | 134188 | 125794 | 131292 | 2900 | 2092 |

**Supplementary Table 2** Read count in different steps of the pipeline.

| Dataset | Name | Taxa | Samples | Min sample depth (reads) | Min sample depth sampleID | Max sample depth (reads) | Max sample depth sampleID | Description |
|---|---|---|---|---|---|---|---|---|
| D1 | ps | 31225 | 82 | 501 | sa81 | 222367 | sa46 | Initial ASV phyloseq object |
| D2 | ps_filtered | 31225 | 81 | 71992 | sa73 | 222367 | sa46 | subset_samples(ps, sampleID != "sa81_BF91") |
| D3 | ps_protists | 27790 | 81 | 31361 | sa17 | 181260 | sa7 | subset_taxa(ps_filtered, Division!=" Metazoa" & Division!="Streptophyta" & Class!="Ascomycota" & Class!=" Basidiomycota") |
| D4 | ps_Sanabria | 31225 | 65 | 71992 | sa73 | 222367 | sa46 | subset_samples(ps_filtered, Description=="Sanabria") |
| D5 | ps_Sanabria_protists | 27790 | 65 | 31361 | sa17 | 181260 | sa7 | subset_taxa(ps_Sanabria, Division!=" Metazoa" & Division!="Streptophyta" & Class!="Ascomycota" & Class!=" Basidiomycota") |
| D6 | ps_parasites | 5925 | 81 | 4658 | sa17 | 49593 | sa14 | subset_taxa(ps_protists, Class==" Opalinata" \| Class==" Labyrinthulomycetes" \| Class==" Oomycota" \| Class=="Ichthyosporea" \| Division=="Apicomplexa" \| Division==" Perkinsea" \| Genus=="Entamoeba" \| Class=="Endomyxa" \| Class==" Endomyxa-Ascetosporea" \| Class==" Chytridiomycota") |

**Supplementary Table 3** Phyloseq datasets generated in this study. Taxa: the final number of ASVs in each dataset. Samples: the number of samples included in the dataset. Min sample depth: the number of reads in the sample with less total reads of the dataset, this is the number of reads to which we rarefy the dataset if needed. Min sample depth sampleID: the name of the sample with less total reads of the dataset. Max sample depth: the number of reads in the sample with more total reads of the dataset. Max sample depth sampleID: the name of the sample with more total reads of the dataset. Description: the phyloseq command for the generation of each dataset.

| samples | Observed | Chao1 | se.chao1 | ACE | se.ACE | Shannon | Simpson | InvSimpson | Fisher |
|---|---|---|---|---|---|---|---|---|---|
| **sa1** | 443 | 465.44 | 10.14009423 | 457.4503763 | 10.49229861 | 4.515093734 | 0.973053776 | 37.11095149 | 73.04764795 |
| **sa2** | 325 | 333.5714286 | 5.749125396 | 330.9590049 | 8.620242695 | 3.563428002 | 0.858284577 | 7.056394963 | 50.5249749 |
| **sa3** | 532 | 551.5238095 | 8.155090023 | 549.7709022 | 11.58872877 | 4.596688438 | 0.973909723 | 38.32845453 | 91.01490448 |
| **sa4** | 311 | 316.6875 | 4.188995322 | 315.7149735 | 8.469177364 | 4.257913875 | 0.968130629 | 31.37809031 | 47.96078098 |
| **sa5** | 305 | 313.25 | 6.362474528 | 308.6952594 | 8.107590566 | 3.492878882 | 0.836769068 | 6.12628983 | 46.86936067 |
| **sa6** | 338 | 339.2173913 | 1.45929306 | 340.3499331 | 9.02701338 | 4.099615812 | 0.953973627 | 21.72667394 | 52.92757426 |
| **sa7** | 520 | 546.6304348 | 9.980385441 | 544.2575129 | 11.50316749 | 4.740186623 | 0.980414127 | 51.05720952 | 88.54664169 |
| **sa8** | 205 | 205 | 0.166259666 | 205.1969364 | 4.894830576 | 3.682316633 | 0.926578995 | 13.62008053 | 29.39766934 |
| **sa9** | 577 | 588.2264151 | 5.328177337 | 589.6173046 | 11.93248085 | 4.690772606 | 0.968887459 | 32.14138033 | 100.3927678 |
| **sa10** | 468 | 479.4545455 | 5.895186452 | 478.6529933 | 10.72944057 | 4.824345624 | 0.983311653 | 59.9220518 | 78.01410445 |
| **sa11** | 241 | 243.5 | 3.161262906 | 242.2809282 | 6.964682505 | 3.601610932 | 0.91211477 | 11.37847616 | 35.52342746 |
| **sa12** | 559 | 571.3333333 | 5.657020199 | 573.5837055 | 11.81866448 | 4.526016022 | 0.955880847 | 22.66589275 | 96.61882151 |
| **sa13** | 366 | 369.5454545 | 2.878730831 | 370.3492395 | 9.129108018 | 4.753539324 | 0.983395705 | 60.22538039 | 58.1710939 |
| **sa14** | 324 | 331.5555556 | 4.983287392 | 329.4729136 | 8.868055449 | 3.525839833 | 0.915455063 | 11.82802943 | 50.34101479 |
| **sa15** | 432 | 441.7307692 | 5.518651315 | 440.9429175 | 10.13599001 | 4.570355694 | 0.977888864 | 45.2260791 | 70.88304436 |
| **sa16** | 562 | 592.1538462 | 11.34223094 | 583.1376027 | 11.85313857 | 4.744170586 | 0.977911786 | 45.27301222 | 97.24571669 |
| **sa17** | 43 | 43 | 0 | 43 | 1.380933287 | 2.794699471 | 0.874074833 | 7.941224348 | 4.907112349 |
| **sa18** | 526 | 542.5 | 7.568183798 | 538.7294657 | 11.44245239 | 4.790785294 | 0.980647398 | 51.6726368 | 89.77903792 |
| **sa19** | 504 | 535.097561 | 11.4663747 | 531.2040526 | 11.38491606 | 4.558247632 | 0.974088549 | 38.59297622 | 85.27735633 |
| **sa20** | 316 | 319.4615385 | 3.106647637 | 319.071568 | 8.312633981 | 4.04331348 | 0.946890775 | 18.82912061 | 48.87376232 |
| **sa21** | 565 | 591.6304348 | 9.980394223 | 587.0037999 | 11.9276677 | 4.701378625 | 0.975828078 | 41.37031406 | 97.87345297 |
| **sa22** | 433 | 448.4 | 8.680293306 | 441.5776556 | 9.919393337 | 4.882435439 | 0.983316308 | 59.93877004 | 71.07929852 |
| **sa23** | 562 | 589.0285714 | 10.73629754 | 580.7262072 | 11.87493378 | 4.526885887 | 0.960110915 | 25.06951436 | 97.24571669 |
| **sa24** | 467 | 487.3125 | 10.51286184 | 476.009822 | 10.61666636 | 4.661532963 | 0.972930433 | 36.94185384 | 77.81421134 |
| **sa25** | 623 | 625.1428571 | 2.096998784 | 625.2712762 | 11.5905908 | 5.324679608 | 0.982947627 | 58.64286436 | 110.1722336 |
| **sa26** | 828 | 847.1666667 | 7.648608633 | 843.3235176 | 14.34213962 | 4.908065436 | 0.93656128 | 15.76324358 | 155.9734667 |
| **sa27** | 907 | 943.5 | 11.28106477 | 938.1319395 | 15.09782231 | 5.066814787 | 0.948365293 | 19.36681834 | 174.5316955 |
| **sa28** | 650 | 663 | 6.821903593 | 657.9409846 | 12.21171052 | 4.918097744 | 0.95001608 | 20.00643414 | 116.0006564 |
| **sa29** | 744 | 753.0357143 | 5.167053887 | 749.6716184 | 13.42522822 | 5.331517356 | 0.981801 | 54.94807324 | 136.782868 |
| **sa30** | 772 | 794.5 | 8.931484242 | 789.0272403 | 13.83053907 | 5.195918732 | 0.972742028 | 36.68651547 | 143.1162474 |

| | | | | | | | | | |
|---|---|---|---|---|---|---|---|---|---|
| **sa31** | 383 | 394 | 6.288468295 | 391.2815771 | 9.595270113 | 4.530604588 | 0.973802334 | 38.17133982 | 61.39904614 |
| **sa32** | 213 | 213 | 0.124706228 | 213.2344206 | 5.629056758 | 3.671549596 | 0.919505062 | 12.42314149 | 30.74180037 |
| **sa33** | 400 | 413.6363636 | 7.259868918 | 409.9434273 | 9.785760354 | 4.578747002 | 0.975533644 | 40.87245286 | 64.65959175 |
| **sa34** | 365 | 369.7142857 | 3.779380057 | 369.0711456 | 9.175746246 | 4.533369158 | 0.974252356 | 38.83850516 | 57.98224581 |
| **sa35** | 485 | 486.1538462 | 1.523855356 | 486.2998304 | 10.0502686 | 4.730307944 | 0.962809299 | 26.88844157 | 81.42780122 |
| **sa36** | 359 | 364.0555556 | 3.777613435 | 363.4836347 | 9.005577704 | 4.387025763 | 0.961808083 | 26.18355078 | 56.85156381 |
| **sa37** | 646 | 652.6666667 | 4.567472884 | 650.0374085 | 12.01776648 | 5.231641958 | 0.978792151 | 47.15235336 | 115.1331323 |
| **sa38** | 490 | 490.8571429 | 1.395265556 | 490.7273544 | 8.404050501 | 5.210017644 | 0.979087997 | 47.81942604 | 82.43736245 |
| **sa39** | 582 | 588.5 | 4.881735254 | 585.3051888 | 10.66778618 | 5.282363041 | 0.980477427 | 51.22275657 | 101.4464135 |
| **sa40** | 703 | 708.5263158 | 3.972981291 | 707.0817563 | 11.7615728 | 5.465827012 | 0.976774378 | 43.05589824 | 127.6261257 |
| **sa41** | 582 | 585.64 | 2.868950217 | 586.2969316 | 11.00320227 | 5.391025557 | 0.989673462 | 96.83787878 | 101.4464135 |
| **sa42** | 388 | 405.6 | 8.142919427 | 403.1799434 | 9.85659111 | 4.170270512 | 0.962031858 | 26.33787064 | 62.35468068 |
| **sa43** | 275 | 276.4285714 | 1.934852552 | 276.2814559 | 6.901463421 | 3.696187019 | 0.875847029 | 8.054579679 | 41.48199727 |
| **sa44** | 504 | 552.4871795 | 16.16758296 | 536.7760757 | 11.37272586 | 4.286532148 | 0.956449698 | 22.96195307 | 85.27735633 |
| **sa45** | 423 | 438.5357143 | 7.5594625 | 436.1392641 | 10.18598794 | 4.435539954 | 0.972216705 | 35.99285103 | 69.12154568 |
| **sa46** | 461 | 466.1333333 | 3.27530155 | 468.7999092 | 10.70887258 | 3.830896548 | 0.917127947 | 12.06679413 | 76.61699974 |
| **sa47** | 520 | 549.9565217 | 10.8662201 | 544.55179 | 11.42010718 | 4.481065302 | 0.969746418 | 33.05393683 | 88.54664169 |
| **sa48** | 338 | 350.047619 | 6.710412333 | 347.8525781 | 8.924637677 | 4.412673209 | 0.973873489 | 38.27529893 | 52.92757426 |
| **sa49** | 274 | 275.25 | 1.621517375 | 275.8154384 | 7.475308004 | 3.744518044 | 0.91948307 | 12.4197482 | 41.30446735 |
| **sa50** | 287 | 293 | 4.699664602 | 290.8643267 | 8.190442476 | 4.042197378 | 0.952188966 | 20.91567399 | 43.62278623 |
| **sa51** | 307 | 310.8823529 | 3.194853392 | 311.817363 | 8.106727375 | 4.381290909 | 0.973791071 | 38.15493507 | 47.23266087 |
| **sa52** | 274 | 275.1538462 | 1.523802514 | 275.6493659 | 7.757436756 | 3.754302204 | 0.937165429 | 15.91480582 | 41.30446735 |
| **sa53** | 329 | 336.65 | 4.920265243 | 335.6860812 | 8.840664018 | 4.129803691 | 0.954901227 | 22.17355221 | 51.26205823 |
| **sa54** | 369 | 389.3076923 | 9.333627973 | 387.1623432 | 9.616071082 | 4.444169516 | 0.974107965 | 38.62191663 | 58.73833032 |
| **sa55** | 359 | 362.4736842 | 2.91045925 | 362.3650853 | 9.255028743 | 3.933503567 | 0.930574404 | 14.40390949 | 56.85156381 |
| **sa56** | 373 | 380.4411765 | 4.36960775 | 382.1246837 | 9.626320063 | 4.117262285 | 0.950978673 | 20.39928462 | 59.49625448 |
| **sa57** | 781 | 788.3076923 | 4.5312327 | 785.8094325 | 13.64684179 | 5.32355488 | 0.982906556 | 58.50196058 | 145.1655584 |
| **sa58** | 535 | 539.5652174 | 3.375376987 | 539.3705962 | 11.17661998 | 5.054993406 | 0.985357488 | 68.29429249 | 91.63413332 |
| **sa59** | 676 | 685.2553191 | 4.780943591 | 685.9766363 | 12.93976459 | 5.170455267 | 0.985902654 | 70.93534024 | 121.6735252 |
| **sa60** | 800 | 815.75 | 7.081756542 | 811.3023668 | 13.99549919 | 5.039559241 | 0.966381622 | 29.74563494 | 149.5133864 |
| **sa61** | 689 | 699.7826087 | 5.316045735 | 700.0058745 | 12.89573983 | 5.005486853 | 0.970768379 | 34.20952996 | 124.5318292 |

| | | | | | | | | | |
|---|---|---|---|---|---|---|---|---|---|
| **sa62** | 777 | 798.5625 | 8.507585957 | 792.914403 | 13.86643786 | 4.953731347 | 0.965952993 | 29.37115725 | 144.2539418 |
| **sa63** | 748 | 759.4473684 | 5.734496261 | 758.9347236 | 12.89621538 | 5.464410211 | 0.983373667 | 60.14555476 | 137.6836952 |
| **sa64** | 613 | 625.4285714 | 6.166615602 | 623.2560667 | 12.19281952 | 5.015769659 | 0.983059101 | 59.02874421 | 108.0299893 |
| **sa65** | 488 | 496 | 5.391138355 | 492.619136 | 10.28410175 | 4.967111826 | 0.984866938 | 66.08047964 | 82.03323802 |
| **sa66** | 673 | 703.3061224 | 10.8030363 | 697.8454631 | 13.00694915 | 4.921470575 | 0.978715229 | 46.98194674 | 121.0159769 |
| **sa67** | 682 | 702.625 | 8.244894319 | 698.5053438 | 13.03668655 | 4.61087986 | 0.956431117 | 22.95216074 | 122.9909431 |
| **sa68** | 341 | 347.5 | 4.724137725 | 345.9809052 | 8.777282135 | 3.806435485 | 0.927358827 | 13.76629746 | 53.48496225 |
| **sa69** | 469 | 486.0322581 | 7.896631284 | 483.1484838 | 10.82369569 | 4.172141574 | 0.956378579 | 22.92451684 | 78.21409953 |
| **sa70** | 673 | 687.0909091 | 6.812131633 | 683.7405426 | 12.68413551 | 5.084268446 | 0.981841001 | 55.06911555 | 121.0159769 |
| **sa71** | 543 | 549.4285714 | 5.468315899 | 544.9389182 | 9.921151914 | 4.905344736 | 0.948008974 | 19.23408846 | 93.28967277 |
| **sa72** | 584 | 585.5555556 | 1.752496538 | 585.5752384 | 11.39138611 | 4.703343698 | 0.935449119 | 15.49165528 | 101.8685161 |
| **sa73** | 434 | 434.6 | 1.059360399 | 434.7994499 | 9.66725998 | 4.339215422 | 0.923363719 | 13.04864983 | 71.27565863 |
| **sa74** | 519 | 520.1538462 | 1.52385986 | 520.1442273 | 10.20750502 | 4.880853208 | 0.961863759 | 26.22177674 | 88.34158103 |
| **sa75** | 731 | 749.3703704 | 8.599985387 | 739.9432212 | 13.28388493 | 5.200778074 | 0.978748593 | 47.05570704 | 133.8643222 |
| **sa76** | 390 | 392 | 2.586103956 | 391.2388593 | 7.711345965 | 4.759062279 | 0.975724173 | 41.19324125 | 62.7377197 |
| **sa77** | 304 | 309.25 | 5.374650717 | 305.53524 | 7.007039269 | 4.395093649 | 0.966378049 | 29.74247396 | 46.68790132 |
| **sa78** | 438 | 439.3636364 | 1.73693875 | 439.3796853 | 9.205483207 | 4.802045372 | 0.974811654 | 39.70089919 | 72.06215628 |
| **sa79** | 722 | 732.7755102 | 5.254877844 | 733.9101387 | 13.34278129 | 4.914831059 | 0.970590694 | 34.00284227 | 131.852022 |
| **sa80** | 303 | 303.8571429 | 1.395190703 | 303.8833971 | 6.74490594 | 4.319122989 | 0.964554902 | 28.21264561 | 46.50656944 |
| **sa82** | 351 | 352.5 | 2.232084903 | 351.8605517 | 6.950224961 | 4.851831641 | 0.981493124 | 54.03396966 | 55.35058368 |

**Supplementary Table 4.** Alpha diversity indices as calculated for each sample by the ASVs abundances.

| Year | Mean Chlorophyll-α (μg/L) | Ecological quality class | Reference |
|---|---|---|---|
| 1973 | 0.7 | High | Margalef et al. 1976 |
| 1974 | 1.9 | High | Margalef et al. 1976 |
| 1987 | 2.46 | High | Pahissa et al. 2015 |
| 1988 | 2.31 | High | Pahissa et al. 2015 |
| 1989 | 2.05 | High | Pahissa et al. 2015 |
| 2013 | 2.16 | High | Pahissa et al. 2015 |
| 2016 | 2.66 | High | This study |

**Supplementary Table 5** Mean chlorophyll *a* values in Sanabria Lake during the last 50 years.

| | | ASVs | | |
|---|---|---|---|---|
| **Effect** | **Df** | **F.Model** | **R^2** | **pvalue** |
| **Site** | 2 | 2.9963 | 0.07135 | *** |
| **Habitat** | 5 | 3.4332 | 0.18625 | *** |
| **Chl_max** | 1 | 2.0148 | 0.02487 | ** |
| **Thermocline** | 3 | 3.1696 | 0.10992 | *** |
| **Sampling site** | 10 | 2.1562 | 0.2355 | *** |
| **Depth** | 5 | 3.1489 | 0.1735 | *** |

**Supplementary Table 6.** Permanova to test for spatial patterns in the protist community structure (p-value ≤0.05 * ; p-value ≤0.01 ** ; p-value <0.001 ***).

| **Grouping** | **R** | **p-value** |
|---|---|---|
| **Site** | 0.6225854 | 0.001 |
| **Habitat** | 0.3950803 | 0.001 |
| **Depth** | 0.4736135 | 0.001 |

**Supplementary Table 7.** ANOSIM analysis calculated upon 999 permutations as a function of sampling site (Sanabria, Laguna, Stream), Habitat (water column, biofilms, sediments) and Depth (Deep, Swallow, Surface).
